# A distributed neural code in the dentate gyrus and in CA1

**DOI:** 10.1101/292953

**Authors:** Fabio Stefanini, Mazen A. Kheirbek, Lyudmila Kushnir, Jessica C. Jimenez, Joshua H. Jennings, Garret D. Stuber, René Hen, Stefano Fusi

**Author notes:** these authors contributed equally to this work (Fabio Stefanini, Mazen A. Kheirbek, Lyudmila Kushnir). Senior and corresponding authors.

## Abstract

The tuning properties of neurons in a given brain region have been traditionally viewed as the underpinnings of computation in neural circuits. However, at the higher levels of processing, specialization is often elusive, instead a mix of sensory, cognitive and behavioural quantities drive neural activity. In such networks, ensembles of neurons, rather than single units with easily interpretable tuning properties, encode behaviourally relevant variables. Here we show that this is the case also in the dentate gyrus and CA1 subregions of the hippocampus. Using calcium imaging in freely moving mice, we decoded the instantaneous position, direction of motion and speed from the activity of hundreds of cells in the hippocampus of mice freely exploring an arena. For the vast majority of neurons in both regions, their response properties were not predictive of their importance for encoding position. Furthermore, we could decode position from populations of cells that were important for decoding direction of motion and vice versa, showing that these quantities are encoded by largely overlapping ensembles as in distributed neural code. Finally, we found that correlated activities had an impact on decoding performance in CA1 but not in dentate gyrus, suggesting different enconding strategies for these areas. Our analysis indicates that classical methods of analysis based on single cell response properties might be insufficient to accurately characterize the neural computation in a given area. In contrast, population analysis may help highlight previously overlooked properties of hippocampal circuits.

## Introduction

The hippocampus has been extensively studied in experiments on navigation and spatial memory. The responses of some of its cells are easily interpretable as these tend to fire only when the animal is at one or more locations in an environment (place cells). However, it is becoming clear that in many brain areas, which include the hippocampus and entorhinal cortex, the neural responses are very diverse^1–4^ and highly variable in time^5–7^. Place cells might respond at single or multiple locations, in an orderly (grid cells) or disorderly way and multiple passes through the same location typically elicit different responses. Part of the diversity can be explained by assuming that each neuron responds non-linearly to multiple variables (mixed selectivity)^1^. Some of these variables may not be monitored in the experiment and hence contribute to what might appear as noise. A neural code based on mixed selectivity is highly distributed because some variables can be reliably decoded only by reading out the activity of a population of neurons. It was recently shown that the mixed selectivity component of the neuronal responses is important in complex cognitive tasks^1, 3^ because it is a signature of the high dimensionality of the neural representations. Place cell discharges are also highly variable^5^ to the extent that the variability, not the spatial tuning alone, can captures changes due to learning in a spatial memory task^7–9^. These recent studies naturally pose the question of how position is encoded within population activity of the hippocampus. To answer this question, we used calcium imaging to record the activity of a large population of neurons in the dentate gyrus (DG), a region of the hippocampus in which the neural responses are highly sparse and diverse^7, 10, 11^, and in CA1, a region that has been extensively studied in relation to spatial navigation using both electrophysiology in rats^12, 13^ and imaging ^6, 14^.

We show that the position of a mouse freely exploring an environment can be decoded from the activity of a few tens of granule cells (GCs) of DG, despite the sparseness and diversity of the code and with an accuracy which is comparable to that of CA1. Using machine learning techniques, we ranked neurons by their contribution to position encoding. We found that trial averaged, single neuron tuning properties are insufficient to predict a neuron’s contribution to position encoding at the population level. Cells that were not spatially tuned according to a statistical test based on spatial information (non-place cells) also contributed to the population code, to the extent that position could be decoded from the ensemble of these untuned cells alone in both areas. We further found that neurons in both DG and CA1 reliably encoded other variables such as the direction and speed of movement. These neurons were not distinct from the neurons that encode position, i.e., the majority of neurons encoded multiple variables and contributed to all of them. We then found that destroying correlated activities among neurons while maintaining their spatial tuning had an impact on decoding performance in CA1 but not in DG. Taken together, these results show that the information encoded at the population level is far richer than at the single cell level and allowed us to uncover the strong degree of stability of DG and CA1 spatial information through the distributed nature of their neural representation.

## Results

We studied the neural code in the DG and in the CA1 area of the hippocampus of freely moving mice. We used miniaturized head-mounted microscopes to perform calcium imaging of granule cells (GCs) in the DG and of pyramidal cells in CA1. To image cell activity patterns we injected a virus encoding the calcium indicator GCaMP6 and implanted a gradient index (GRIN) lens for chronic imaging (Fig. 1). Four weeks after surgery, we imaged cellular activity while mice foraged for sucrose pellets in an open field arena. We then used a recently developed algorithm for reliably extracting the GCaMP signals from the raw videos, CNMF-e^16^(Fig. 1d, e). This algorithm separates local background signals due to changes in fluorescence in the neuropil from the signals due to calcium concentration changes in individual cells and was particularly effective in identifying signal sources in our granule cells data without introducing spurious distortions or correlations among cells due to artifacts. We recorded a total of 1109 DG cells across 3 animals, among which 352 (32%) were significantly tuned to position and a total of 863 CA1 cells across 3 animals, among which 38 (4%) were significantly tuned to position (see Methods and Fig. S2).

**Figure 1.**
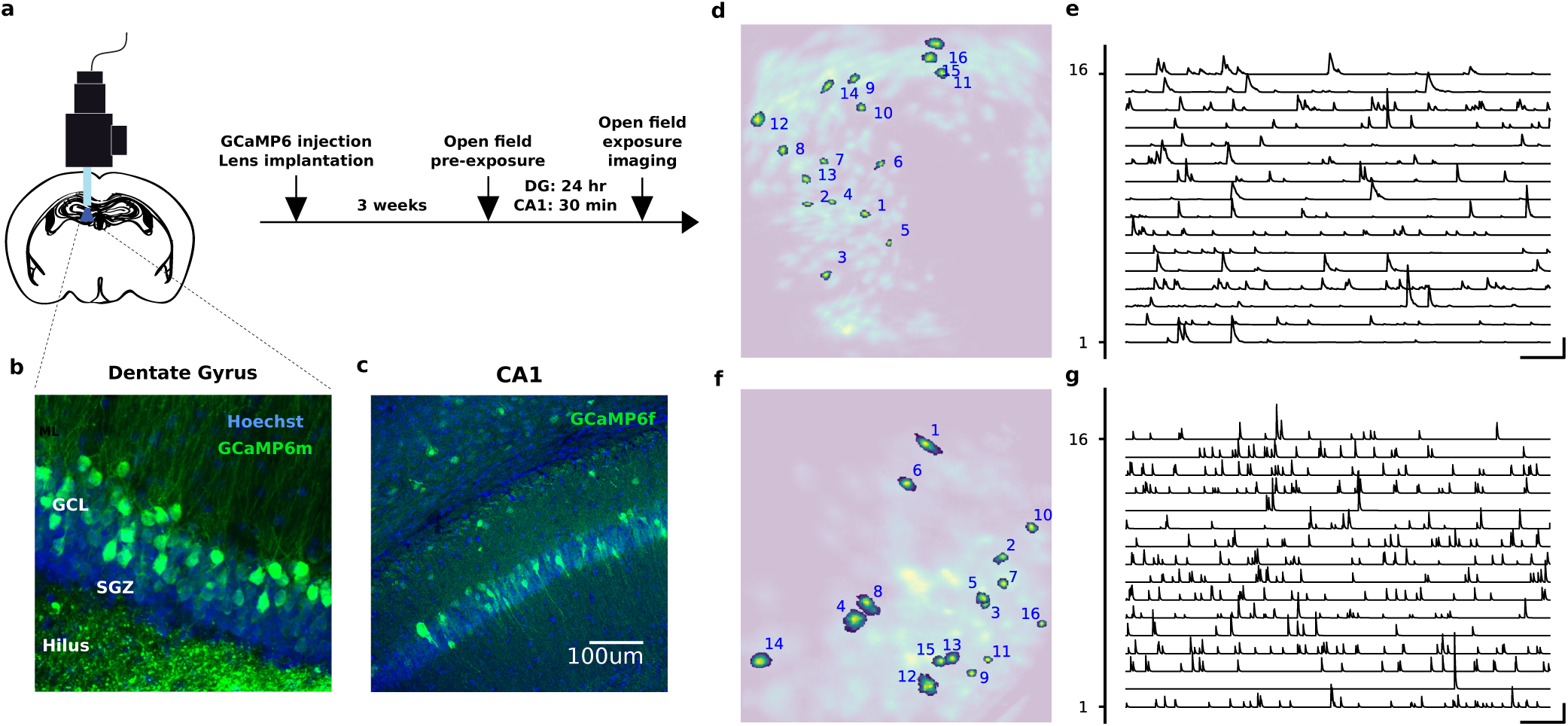
**a**) Experiment protocol. Mice were anesthetized with isoflurane and placed in a stereotactic apparatus. DG mice were then injected in the dorsal DG with a virus encoding GCaMP6m. Ca1 mice were injected with GCaMP6f. Mice were then implanted with a GRIN lens and a baseplate was attached to the skull at the optimal imaging places. Three weeks after surgery they were checked for GCaMP expression with a miniaturized microscope (Inscopix, Palo Alto, CA) and procedures previously described^15^. Recording site. The imaging plane was later assessed through histology. **b**) DG recording site. GCL, granule cell layer; SGZ, subgranular zone. **c**) CA1 recording site. **d-g**) Automated signal extraction using CNMF-e^16^. The algorithm identifies the spatial (left) and temporal (right) components of the signal sources, i.e., putative cells. It uses a generative model of Calcium traces and non-negative matrix factorization to separate actual signal sources from the background due to diffused neuropil fluorescence. The extracted spatial components are displayed in **d** (DG) and **f** (CA1), where a few representative ones are highlighted. The corresponding temporal profiles are shown on panels **e** (DG) and **g** (CA1). The traces are obtained by convolving the inferred Calcium events with a temporal profile resembling the Calcium indicator dynamics (see Fig. S1). Scale bars are 10 min and 10 sd.

The low fraction of place cells in CA1 seems to be in contrast with reports from previous studies in CA1^6, 17–19^. However modern tools for source extraction from calcium imaging can detect cells with very low activity, which are largely underestimated in electrophysiological recordings as other recent studies have also suggested^20, 21^. If one excludes from the analysis these low firing rate cells, the fraction of place cells can be significantly higher (see Fig. S3).

The first step of our analysis was to assess whether the position of the animal is encoded in the recorded neural activity during mobility. We therefore removed all the time bins in which the animal was slower than 2 cm/s for a period longer than 1 s, after confirming by visual inspection that this procedure would exclude moments of immobility. To decode position, we discretized the *x* and *y* coordinates of the animal by dividing the arena into 64 regions (8 by 8 grid) (50 cm square arena for DG mice, 50 × 28 cm for CA1 mice). We then trained a battery of linear classifiers for each pair of discrete locations. Each session was divided into 10 one-minute long intervals, 9 of which were used to train the classifiers and the remaining ones to test them (10-fold cross validation). We used a majority rule^22^ to combine the outputs of the linear classifiers as an instantaneous estimate of the animal’s location, using the center of the selected location as the decoded position.

In both areas, the median decoding error was comparable to the animal size, revealing for the first time that instantaneous position can be decoded from DG GCs population activity (Fig. 2). Our analysis of CA1 show comparable decoding accuracy between DG and CA1 after correcting for number of cells (Fig. 2c and Fig. S4) and is slightly higher than the one observed in previous studies in CA1^6^. Different decoding strategies, such as using linear decoders and Bayesian decoders, as well as decoding from raw Calcium traces or events, produced similar results (see Fig. S6). The decoding error was found to weakly correlate to the speed of movement (see Fig. S7). To our knowledge, this is the first time that decoding of position from populations of DG cells has been reported.

**Figure 2.**
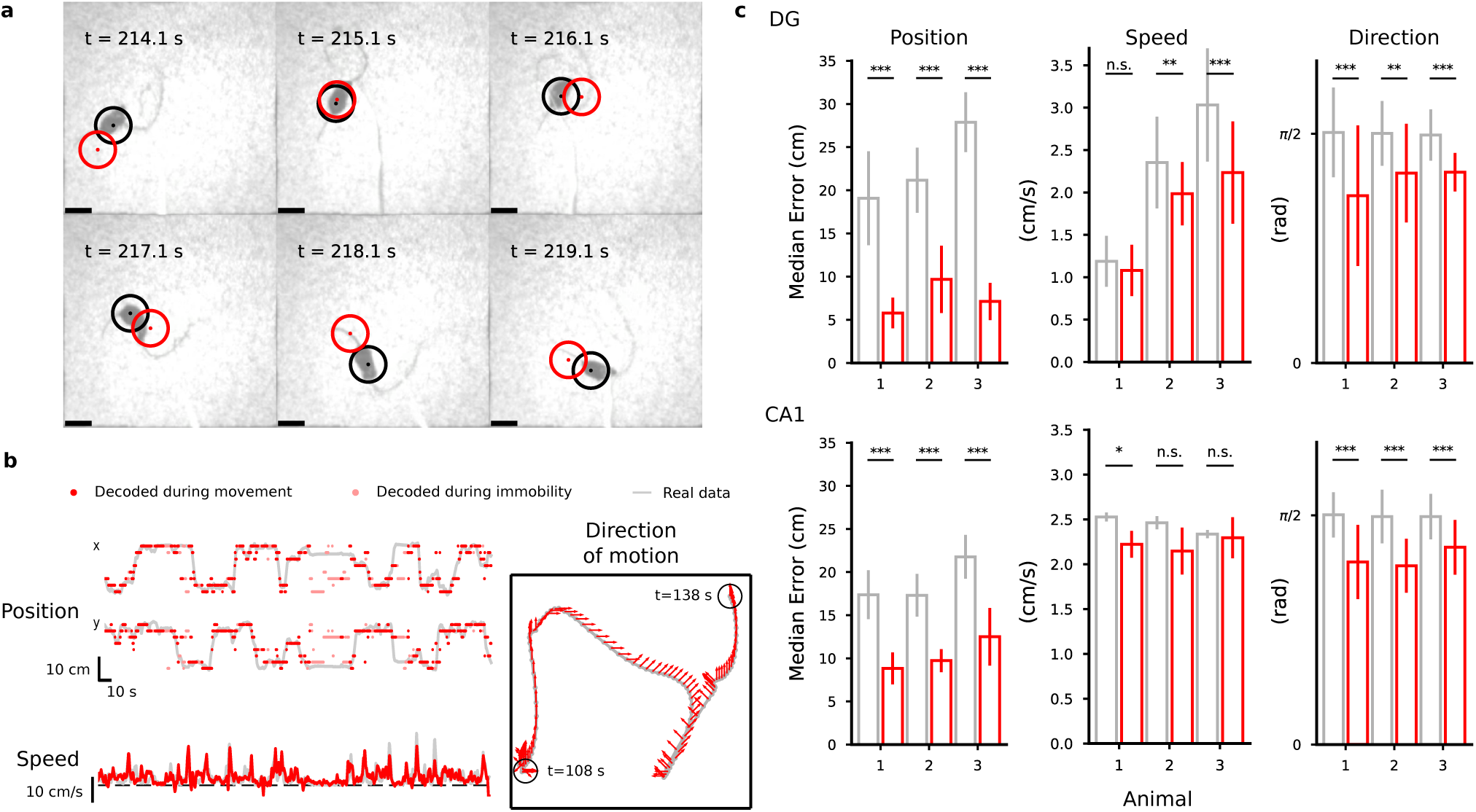
Decoding position, speed and direction of motion. **a**, **b**) Decoding results for a representative DG mouse. See Supplemental Material online for a decoding video. **a**) Selected frames of a video showing the arena and a DG animal from above. The black circle represents the mouse’s actual position and red circle is the decoded position, obtained with a Bayesian decoder that reads the activity of 317 DG cells. Neural activity has been pre-processed to identify putative calcium events as explained in the Methods. **b**) Examples of decoding position, speed and direction of motion (DG representative mouse). Grey lines correspond to the real values of position and speed variables in the top left and lower left panels respectively while the red dots correspond to their decoded values. The time bins marked in light red in for position and direction of movement correspond to moments of immobility that have been excluded from the training data. The grey line in the right panel corresponds to the position of the mouse and the red arrows correspond to the decoded direction of motion in a 30 s time window. **c**) Decoding accuracy (top: DG; bottom: CA1). The decoding error for position and head direction is computed as the median distance between the decoded value in each time bin and the actual value of the decoded variable in the test data. For the direction of motion, the smallest angle between the decoded and the actual value is considered. The red vertical bars corresponding to the mean over the 10-fold cross-validation (error bars correspond to st. dev.). Grey: chance error obtained by decoding from shuffled data in a way that preserves the autocorrelations in the data (see Methods and Fig. S5). Number of cells: 483, 309, 317 in DG mice; 371, 286, 206 in CA1 mice. See also Fig. S4-S7, S12.

We could also decode the direction of motion of the animal in both regions and its speed only in DG. Speed was weakly correlated with the overall level of activity in DG and we could decode it in two animals out of three using linear regression (Fig. 2b, c). To decode the direction of motion we divided the full range of possible directions into 8 angular bins and labeled time bins according to the instantaneous discrete direction of motion of the mouse (see Methods). To our knowledge, this is also the first time that decoding of direction and speed of motion from populations of DG and CA1 cells has been reported although direction tuning has been previously observed in CA1 pyramidal cells in rats^23^. We did not find differences in decoding performance for direction of motion between the DG and CA1 areas (Fig. 2c).

To better characterize the neural code, we tried to determine what features of the response properties of individual neurons are important for encoding the variables we could decode. It is important to realize that these response properties could be dissociated from the contribution of a cell to the accuracy of a decoder that reads out a population of neurons. For example, there could be neurons that are only weakly selective to position and so individually would not pass a statistical test for spatial tuning. However, when combined with other neurons, they can still contribute to position encoding. Alternatively, the decoder might assign a large weight to neurons that are weakly selective. This situation can be illustrated with the intentionally extreme case of Fig. 3, in which we show how the responses of individual neurons can be dissociated from their importance for the decoder. A simulated animal visits two locations of the arena multiple times. The activity of two hypothetical neurons is represented in the activity space (Fig. 3b), with the horizontal and vertical axes representing the activity of the first and the second neuron respectively. At each pass through each location the two neurons have different activity due to other variables that might also be encoded, e.g., the direction of movement, the speed of the animal, or other variables that are not under control in the experiment. Each point in the activity plot represents the activity of the neurons in a single pass. The responses of neuron 2 to the two different locations have the same distribution (Fig. 3b,c). A cell with such response properties is untuned to space (a non-place cell) and therefore it is typically considered unimportant for encoding position. However, a linear decoder trained to decode the position of the animal can make use of the untuned neuron because of the correlations between the activities of the two neurons. While the activity of neuron 1 is only partially predictive of the animal’s location (the distributions partially overlap), by reading out neuron 2 together with neuron 1 it is possible to decode position with no errors using a linear decoder. In such situation, the linear decoder would assign equal weights to the two neurons, as shown in Fig. 3c.

**Figure 3.**
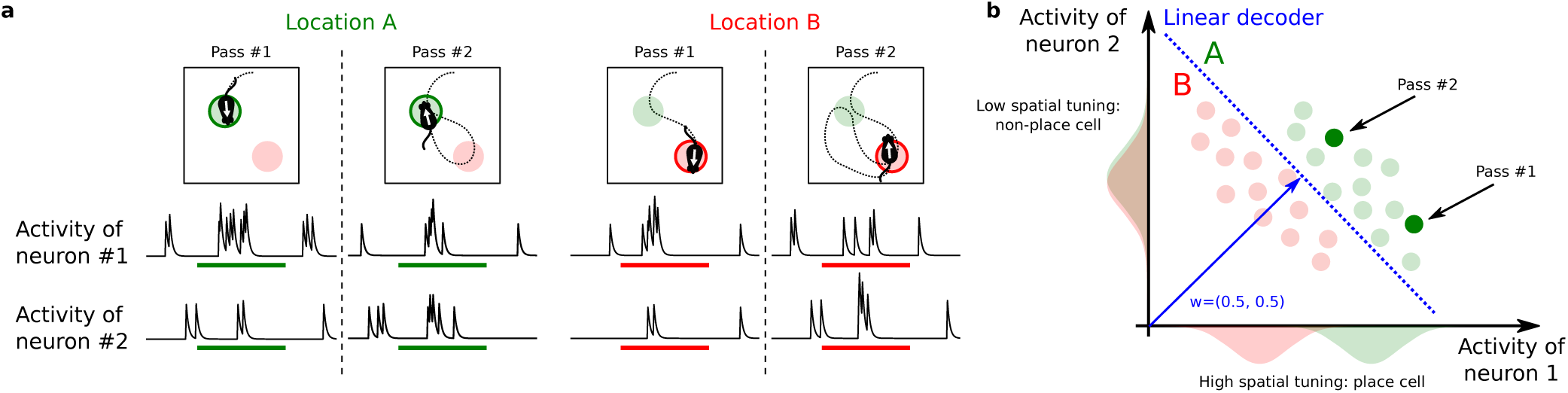
The contribution of untuned cells for encoding position. We show an extreme situation in which one simulated neuron has the same activity distribution when the animal is in two different locations of the arena. Hence the neuron is not selective to position. Nevertheless, for a decoder this neuron can be as important as other selective neurons due to its contribution to the population coding. **a**) Activity of two simulated neurons as a function of time. Top: The simulated animal visits the same discrete location twice (location A in green, location B in red). Bottom: Simulated traces around the time of passage through each location. Different responses for the two neurons are elicited by different experiences, for example due to the different direction of motion. **b**) Example of how place cells and non place-cells can be equally important for encoding the position of the animal. In the scatter plot, the x-axis represents the average activity of the first neuron during one pass and the y-axis is the activity of the second neuron. Each point in the space represents an average population response in a single pass. Their responses are typically highly variable and are scattered around their mean values. The two neurons in the example have very different activity profiles: the first has a strong spatial tuning (place cell) while the second has only a weak tuning. The distributions of their activities in each location, reported along the axis, only partially (neuron 1, place cells) or almost completely overlap (neuron 2). Despite this variability in the single neuron responses, the neural representations at the population level are well separated, making it possible for a linear decoder (blue dashed line) to discriminate them with high accuracy. The resulting decoder’s weight vector has two equal components corresponding to the importance of the two neurons in encoding position. In this example both neurons are important for encoding position despite their very different tuning properties.

In the real data, there might be a spectrum of different situations that are less extreme than the one illustrated in Fig. 3 in which a decoder can take advantage of weakly tuned cells. Cells like the untuned one in Fig. 3 or weakly tuned cells can “cooperate” with more tuned cells to encode more precisely a variable like position. This is a situation similar to the one of Fig. 3, in which the correlations between the activities of different neurons would be important. However, there might also be weakly tuned cells that are uncorrelated, but when combined together would contribute to the accuracy of a decoder. In both cases, the decoder can use the weakly tuned cells to improve its accuracy. Analogously a downstream neuron can in principle harness the activity of weakly tuned neurons to readout the animal’s position.

In our analysis we took the perspective of such a readout neuron and analyzed the weights assigned to cells by our decoder to determine the importance of input neurons in a population for encoding position. The procedure we adopted was to first train the position decoder on each pair of locations and then to combine the resulting weights to obtain a single importance index (*ω*) for each cell (see Methods). Similar methods are used to assess the importance of individual features in a feature space^24, 25^ and have been recently used to identify important synapses in learning models^26^. We then ranked the neurons according to this importance index and estimated the decoding accuracy for populations of 50 neurons (Fig. 4a) to assess the validity of our approach. The 50 neurons with the largest importance index indeed performed significantly better than the worst 50 neurons, though position could be decoded above chance level even from the worst neurons. The accuracy decreases progressively between the performance for the best and for the worst neurons, validating the method for ranking the neurons on the basis of the importance index. We also controlled that the ranking was stable within the session (see Fig. S8).

**Figure 4.**
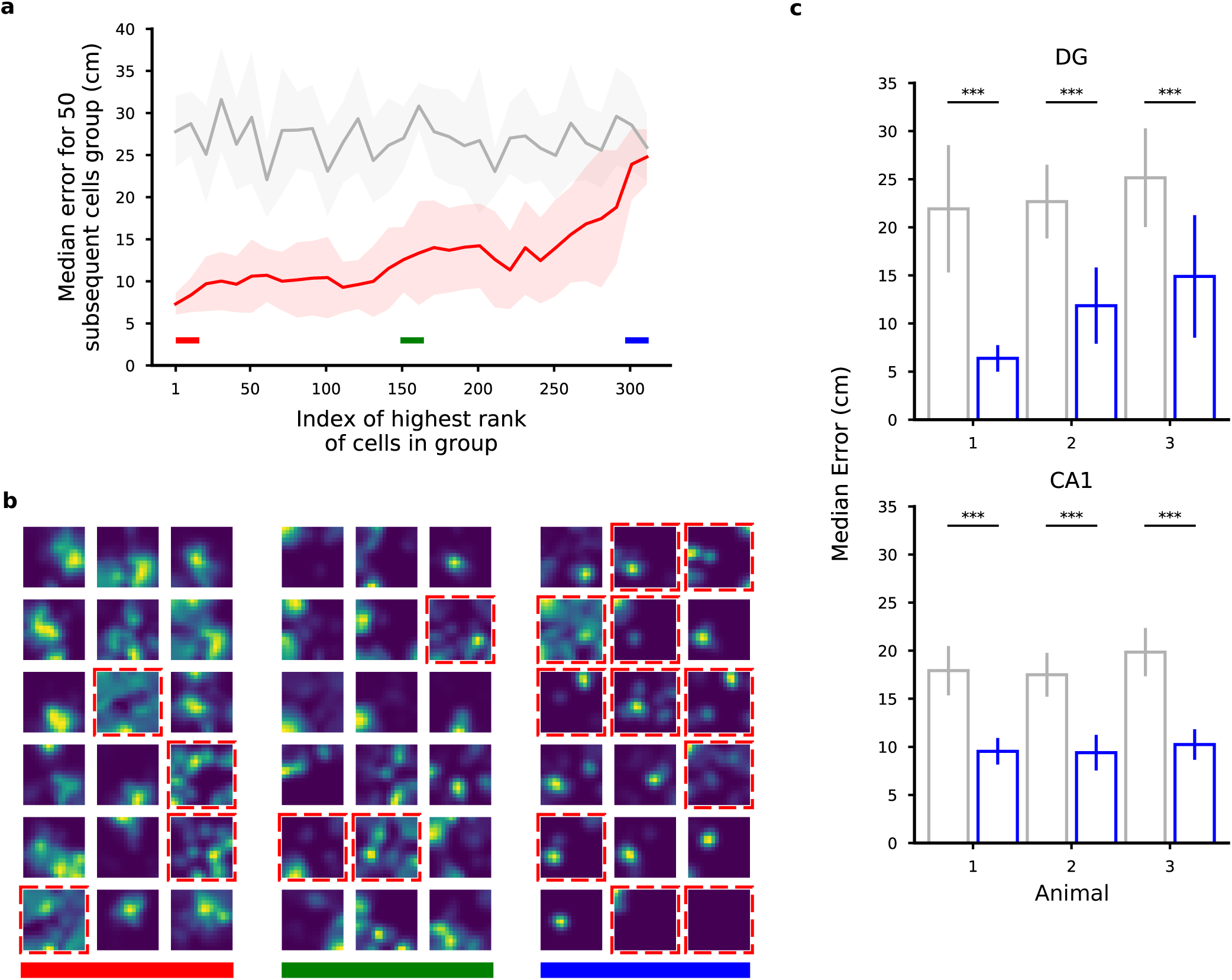
Ranking neurons according to their contribution to the decoding accuracy for position. **a**) Validation of the importance index. In this figure we show the median error for various selections of 50 DG cells from a representative animal ranked by their importance index as obtained using the decoder’s weight. Each point in the plot is aligned to the rank of the first cell in the selection (for example, the first dot corresponds to the selection of the first 50 cells from index 1 to index 50; the shaded region represents the standard error for the 10-fold cross-validation). Grey: chance level and standard error. As expected, the median error for the population of the 50 top ranked (best) cells is much smaller than the median error for the worst last (worst) 50 ones. **b**) Spatial tuning maps for groups of 18 cells ordered by importance index. Same cells as in **a**. We ranked the cells using the importance index for position (see Methods). The three groups of best, mid and worst cells are highlighted with the color bands in **a** for reference. The maps are normalized to the peak rate in each map. Dashed red borders indicate cells that don’t pass the criteria for place-cells using a commonly used statistical test for tuning (see Methods). Even among the most important cells there appear some non place-cells (and vice versa). Similarly, some place cells appear in the group of cells with medium and low importance. The small fields in the group of low importance cells are due to significantly lower activities (see also Fig. 5). **c**) The position of both DG and CA1 animals can be decoded from the activity of the non-place cells with a performance significantly higher than chance (Mann-Whitney U test, ***p<0.001). Number of cells: 451, 208, 98 in DG mice; 350, 277, 198 in CA1 mice. See also Fig. S2, S3, S8.

The observation that most neurons could contribute to the decoding of position indicates that the neural code is highly distributed. Indeed, the importance index is rather similar for most of the cells. If the importance index was a measure of “wealth” and cells were households, the hippocampus would be the country with the lowest inequality in the world (see Fig. S13 in which we estimated the Gini coefficient for the importance index).

Not too surprisingly, one important feature of an individual neuron is its average activity, which is strongly correlated with the importance index and hence to the overall ability to encode position (see also Fig. 5a, b). However, inspection of the firing fields of Fig 4b indicated there were no other obvious properties that predicted whether a neuron is important or not in both DG and CA1 neuron populations. We then identified which neurons where spatially tuned and called them place cells if the spatial information contained in their activity was statistically significant (see Methods for details). The difference between the spatial information for the recorded activity and the spatial information obtained for shuffled data, properly normalized, is what we defined as significance of spatial information (SSI). It is indeed a measure used to assess whether a cell is a place cell or not relative to a null distribution^18, 27, 28^.

**Figure 5.**
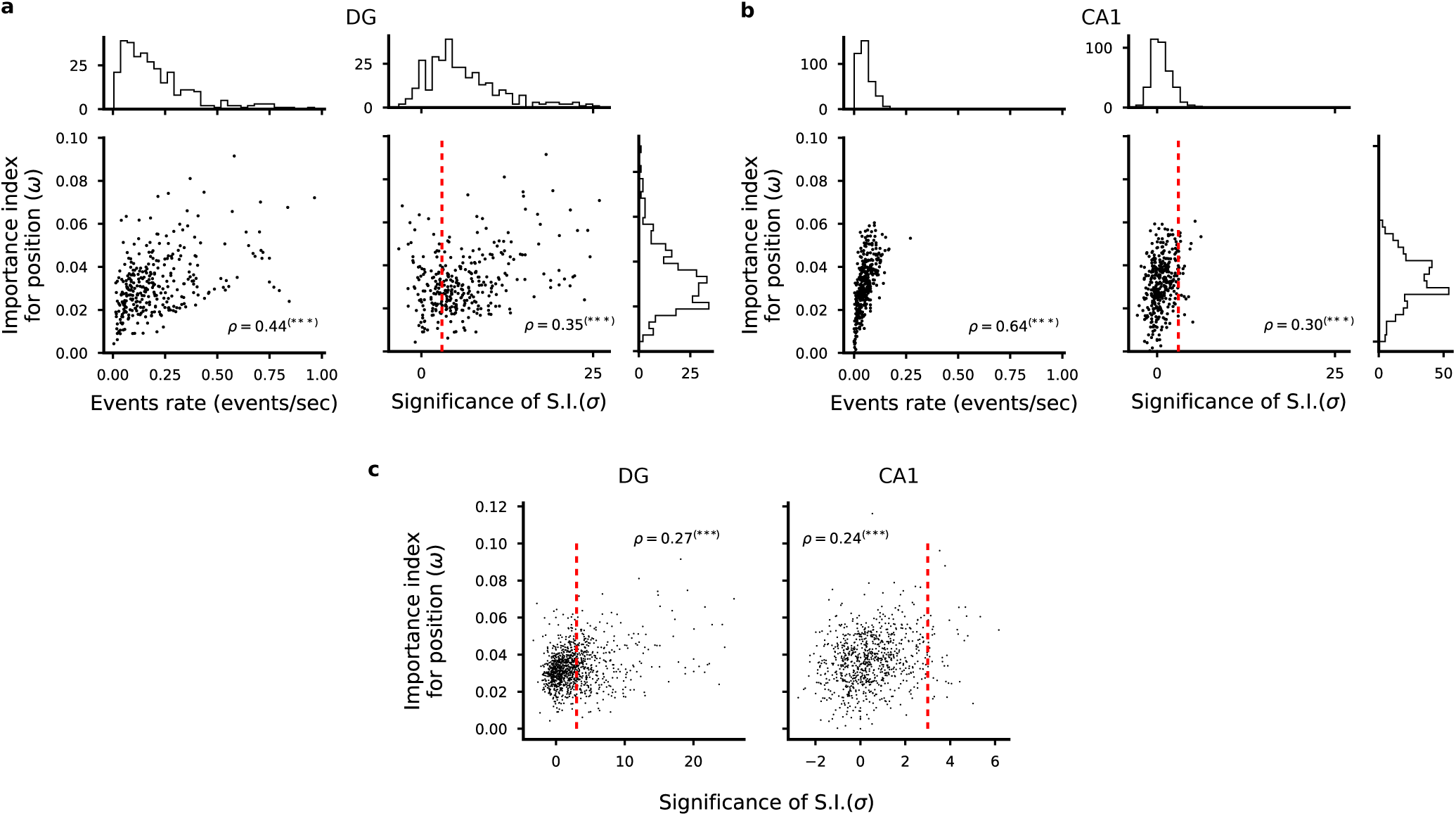
Correlation between importance index and spatial information. **a-b**) Left: Scatter plot of importance index and overall cell activity for each cell in one representative animal. As expected, we found a strong correlation between these quantities because it is unlikely that a weakly active cell can contribute to decoding. Right: Scatter plot of importance index and statistical significance of spatial information with respect to independent random temporal shuffling of each cell’s identified calcium events. DG cells in **a**, CA1 cells in **b**. Each dot corresponds to one cell in one representative animal. Pearson’s correlation factor *ρ* between the plotted quantities are reported (***p<0.001). Significant correlations are found between the analysed quantities but single cell statistics only partially capture the information available at the population level. For each quantity, overall histograms are reported on the side of the plot. The dashed red line corresponds to a value of a threshold of 3 used to define place cells (see Methods). **c**) Same plots as in **a** and **b** but for all cells identified in DG (left) and CA1 (right). See also Fig. S9, S13.

From Fig. 4b it is clear that there are non-place cells that have a large importance index. The animal position can be decoded from these cells alone in both DG and CA1 (Fig. 4c). This indicates that the activity of the non-place cells population contains some spatial information. However, because of noise and limited data, the activity of these cells does not pass the statistical test that we adopted to characterize place cells.

While the SSI is a property of the single cell, the importance index depends on the contribution of a cell to the population code. We thus analyzed the relation between each cell’s SSI and its importance index. Although there is not a one to one correspondence between SSI and importance index, the two quantities are correlated (Fig. 5a, b, c) indicating that some individual response properties are at least partially informative about the importance of a cell in encoding position. To compute the SSI one has to compute the spatial information and subtract a baseline obtained by shuffling the activity^11, 18^. The spatial information without the baseline subtraction, which is sometimes used as a measure of the tuning of the cells^27^, is actually negatively correlated with the importance index (see Fig. S9). This is a reflection of the sampling bias problem that affects cells with low activity^28^.

We performed a similar analysis of importance for the direction of movement. In Fig. 6 we show that we can rank the cells according to their contribution to decoding (Fig. 6a) and that the important cells are highly heterogeneous in their direction tuning (Fig. 6c). Considering all recorded cells, we also found that a cell’s activity correlated with the importance index for direction of movement in DG (Fig. 6c) but not in CA1 (Fig. 6d). We defined the significance of directional information (SDI) in a similar way to the SSI by comparing the mutual information between direction of motion and a cell’s activity to a distribution obtained by shuffling the cell’s calcium events in time. The importance index and this directional information were correlated both in DG (Fig. 6c) and in CA1 (Fig. 6d).

**Figure 6.**
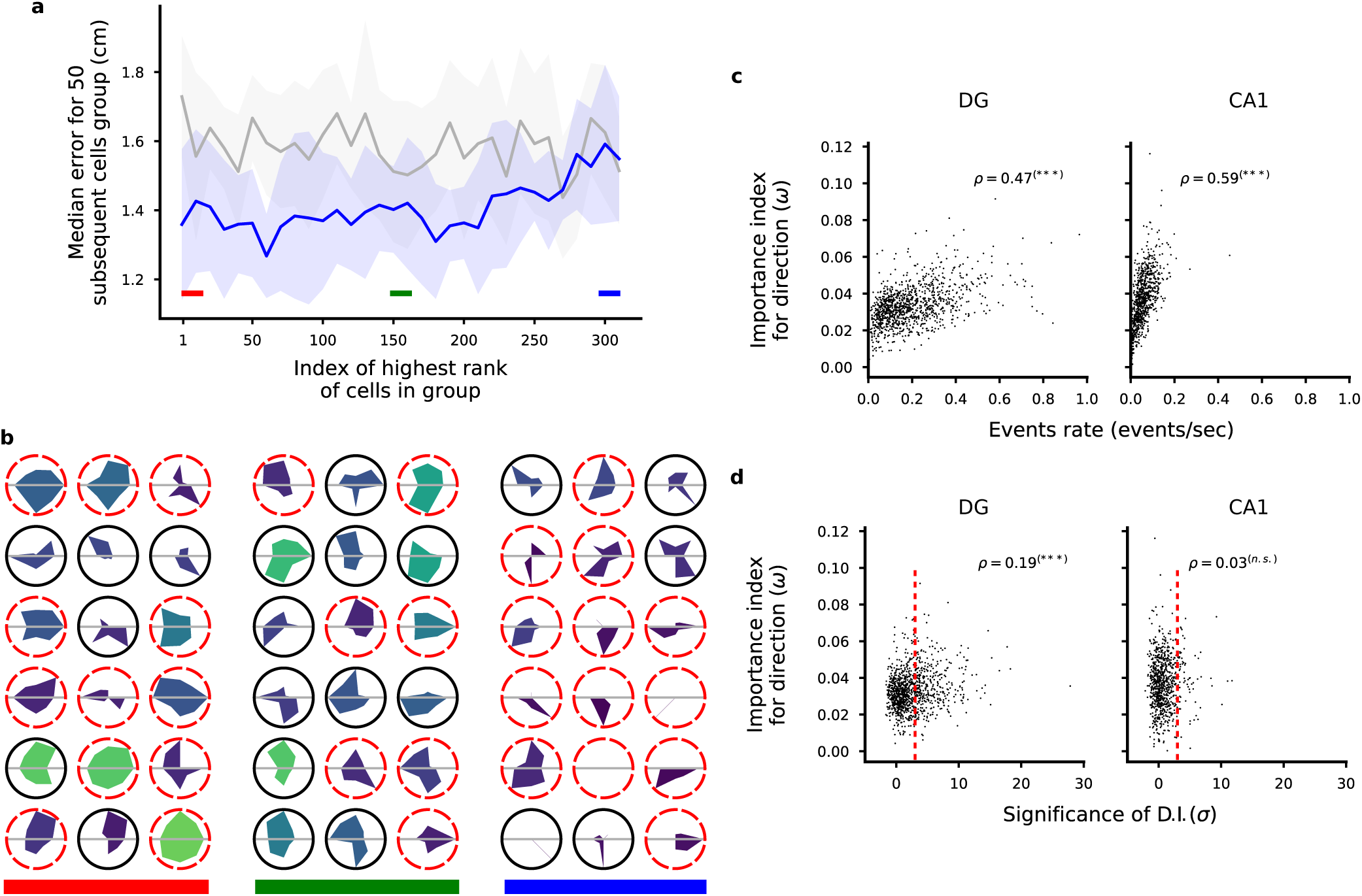
Ranking neurons according to their contribution to the decoding accuracy for head direction. **a**) Validation of the importance index as in Fig. 4a but we ranked the cells according to the importance index for decoding direction of motion (see Methods). **b**) Tuning maps as in Fig. 4b. Here we show the tuning for direction of motion of single cells as polar tuning maps for groups of 18 cells ordered by importance index. The area colour represents the overall activity of the cell throughout the trial. Dashed red borders indicate cells that don’t pass the criteria for significant direction tuning using a commonly used statistical test (see Methods). As in the case of position tuning, some untuned cells appear among the most important cells and highly tuned cells appear among the least important. **c**) Scatter plots of cell activity and importance for position decoding for all identified cells in DG (left) and CA1 (right). Pearson correlation factor *ρ* between the plotted quantities are reported (***p<0.001). **d**) Same as in **c** but for importance index for direction and significance of direction information.

All these analyses indicate that single neuron properties are only partially predictive of the importance of a cell for decoding. Moreover, the importance is not an intrinsic property of an individual cell because it clearly changes depending on which other cells are part of the population of neurons that are used by the decoder. This is clearly illustrated in 3, in which the untuned cell is important when combined together with the cell represented on the horizontal axis, but it would be useless if combined together with another untuned cell, or with an uncorrelated tuned cell.

Since we could decode at least two variables from the neural activities, we were wondering whether there is some form of specialization in which segregated groups of neurons encode different variables. In Fig. 7 we report the importance index for the direction of movement versus the importance index for position (Fig. 7a). The situation in which different variables would be encoded by segregated populations of neurons would predict a negative correlation between these two importance indices: cells with large importance index for position should have a small importance for the direction of movement, and vice versa. For both the regions we analyzed, we found a positive correlation between the two quantities, with a higher correlation in CA1, suggesting that neurons that are important for encoding one variable are also important for encoding the other. This is partially explained by the fact that for both position and direction of movement the most active cells tend to be the most important ones. However, when we regressed out the components explained by the activity, we still found a positive correlation between the importance indexes of the two variables (Fig. 7a). In addition, this could not be explained by correlation between direction of movement and position (Fig. S11).

**Figure 7.**
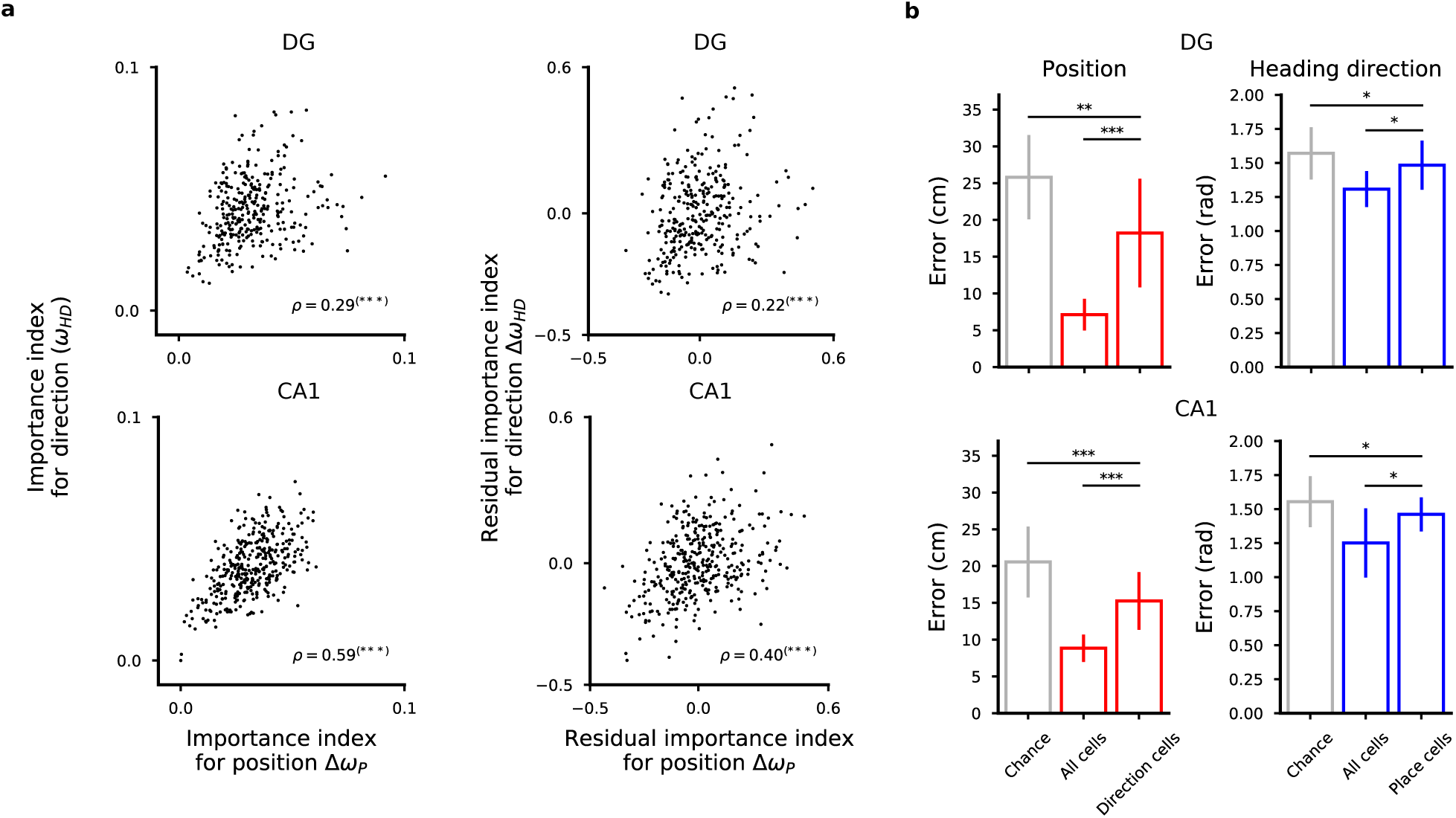
The representations for space and direction of motion are distributed in DG cells and CA1 cells. **a**) Left: Scatter plots of importance index for position and direction of motion (top: DG cells in one representative mouse; bottom: CA1 cells). Each dot corresponds to one cell for which we computed the importance index for the variables we decoded. Pearson’s correlation values *ρ* are reported (***p<0.001). Right: same as in left but the component of the correlation due to the correlation between importance index and cell activity has been removed from the data. Residuals from linear regression are considered for both quantities. Also the residuals show a positive correlation. **c**, **f**) Even the most important cells for encoding one variable carry information about the other variable. We show the decoding performance of position (left) and direction of motion (right) using the most important cells for direction and position (left and right plots respectively). See also Fig. S7.

We then focused on those cells that had a high importance for one variable and not for the other as candidate specialized cells. However, we could decode position from the most important cells for encoding direction of motion and vice versa, showing that even the most important cells for one variable carry information about the other variable in both regions (Fig. 7). We conclude that in both CA1 and DG, neurons have mixed selectivity to the variables we decoded, in line with recent studies both in CA1^18^ and in EC^4^ as well as in the cortex^1, 3, 29^.

So far, we have shown that the code that is used to represent position is distributed, i.e., all active cells contribute to some extent to the population code. We therefore sought to see if correlations between the activities of different neurons contribute to the decoding performance. In general, it is possible that noise correlations are beneficial, detrimental or irrelevant for the neural code^30–32^. However, our intuition is that a large portion of the noise variance can be explained by the fact that neurons encode multiple variables (see Discussion). For example, the different points that encode the same position in Fig. 3 might correspond to visitations in which the head direction and/or the speed were different. In this case, destroying the correlations would result in a decrease or no change in decoding performance (Fig. 8a). We devised a procedure to shuffle the data in a way that destroys the correlations across neurons while maintaining the spatial tuning of each cell (see Methods). We then studied the effect of this procedure on the decoding accuracy. For example, consider the problem of decoding position. At each pass through a location, we randomly picked the activity of a cell from the pool of recordings corresponding to that location and that cell (Fig. 8b). We then corrected for the different time spent in each pass at the same location and repeated the procedure for all cells independently. By using this procedure we effectively destroyed the correlations among neurons because, after this manipulation, each cell’s activity was independent from the others. However, by restricting the manipulation to each discrete location, we did not alter the spatial tuning of the cells (Fig. 8c) and the correlations among neurons induced by their tuning profiles. Finally, by comparing the performance of the decoder on the new data we could assess the contribution of the correlations to decoding, which is a direct test of the presence of a structure in the neural representations that is beneficial for representing information^30, 33–35^. In 4 of the 6 analyzed animals we found that the decoding error increased when correlations were destroyed through the shuffling procedure, revealing the importance of orchestrated activity within the population (Fig. 8d). The effect was very consistent in CA1 neurons where performance was reduced by about 20%, whereas almost no effect was observed in DG.

**Figure 8.**
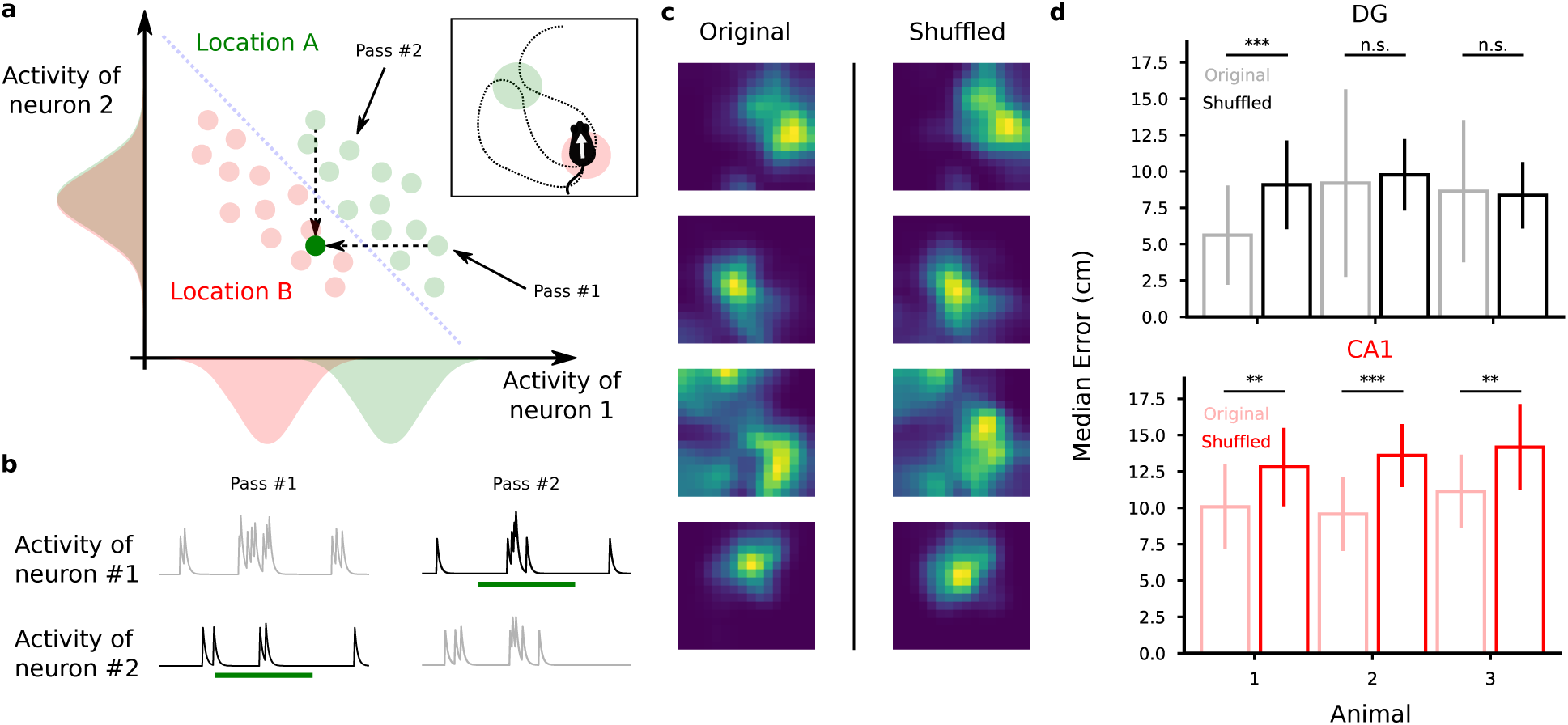
Destroying correlations impacts decoding performance in CA1 but not in DG. **a**) Procedure to test the presence of correlations between cells. We record neural activity during multiple passes through location A (green). Then for we generate a new recording by randomly choosing one of the activities recorded in that location for each cell independently. The green dot below the decoder’s discrimination line in the activity plot corresponds to the newly generated activity. We repeat this for all the passes through each location and for each cell independently, therefore destroying the correlations between cells, if any. In the extreme case depicted in the cartoon, this procedure will introduce errors in decoding position since the generated activity will be classified as the wrong location.**b** Cartoon activity traces for the two correlated neurons during the two passes through the same location. As described in **a**, we destroyed correlations by choosing for each neuron the activity during one of the passes through that location and combined them to generate a new activity pattern corresponding to that location. In this example, we chose pass 2 for neuron 1 and pass 1 for neuron 2. **c** Spatial tuning maps of four representative cells before and after the suffling procedure to destroy correlations. The spatial tuning of the cells remain unaltered after the procedure (mean st. dev.; Mann-Withney U test, ***p<0.001, **p<0.01). **d** Decoding performance before (light colors) and after destroying correlations through shuffling (full colors). Top: DG animals. Bottom: CA1 animals. See also Fig. S14-S15.

## Discussion

Neurons in the DG and in CA1 have rather diverse response properties and often the responses are not easily interpretable^10, 11^. Despite this seemingly disorganized neural code, it is possible to decode from the activity of a population of neurons the position, the speed and the direction of motion of the animal. Neurons respond to mixtures of the decoded variables as observed in other high cognitive brain areas^1, 3^. The information about these variables is highly distributed across neurons to the point that the responses of individual neurons are only weakly predictive of their contribution to the neural code. It is therefore crucial to consider neurons in one area as part of an ensemble to assess their importance for processing and transferring information about a particular variable.

One implication of such distributed neural code is that it can be misleading to characterize the function of a brain area based only on the statistics of individual neuron properties. In the specific case of position encoding, for instance, it is not possible to conclude to what extent the position of the animal is encoded in a brain area only by analyzing the tuning of individual cells to space. Indeed, populations of cells whose activities do not pass a selectivity criterion for space encoding, for example through an information theoretical approach, may still encode position via the ensemble activity patterns, as we showed by decoding position and direction of motion from untuned cells in both DG and CA1 regions of the hippocampus.

The population coding rescues the ability of these areas to encode position despite the sparsity of its activity and the variability of its representations. Here we show that indeed even few tens of cells encode position with high precision in both analyzed areas. Furthermore, the decoding was accurate even when model training and model test periods were separated by up to 18 minutes, indicating that at the population level the representations were stable, despite the elevated variability of individual cells (see Fig. S12).

Our findings are in line with studies suggesting that session averaged, single cells statistics falls short in describing the activities of hippocampal cells. For example, while place fields are widely used to analyze DG activities in remapping studies^10^, it is only when sub-second network discharge correlations are taken into account in the analysis that memory discrimination signals can be revealed^7^.

Destroying correlations among neurons did not have a strong impact in decoding performance in DG neurons but it consistently reduced decoding performance in CA1 data. We conclude that the traditional model of point representations in the neural space is inappropriate for spatial representations carried by neurons especially in CA1. Whether neural correlations are used in the population code is a long-standing question. In the data, it has been shown in the past that the pair-wise correlations accounted for about 10% of the information contained in neural activities^33, 36, 37^ while using models that exploit higher-order correlations can recover about 20% of information related to the stimulus in a population of retinal ganglion cells^34^. Here we show the same impact of correlations in CA1 but not in DG. This could reflect the different type of input patterns impinging on these regions.

These observations indicate that the correlations that we are destroying should be considered as signal correlations rather than noise correlations, at least in CA1. The variability across visitations can probably be explained by the fact that neurons encode multiple variables in a consistent way and may induce the observed neural correlations. This situation would be similar to the one discussed in Figure 3 (e.g., pass 1 and 2 would correspond to two visitations of location A with a different direction of motion), i.e., the disruption of the correlations decreases the performance of the decoder. In Fig. S15 we show in simulations that this is indeed the case. We considered a model in which the neural activity depends on multiple variables (e.g. the position of the animal, the direction of motion etc). Each variable has a discrete set of different values and every set of values of the encoded variables corresponds to one specific condition. We considered two different scenarios in which we constructed neural representations that are inspired by previous theoretical and experimental studies. More specifically, we considered 1) unstructured representations in which different conditions are represented by different random vectors in the activity space 2) structured representations in which the geometrical structure is the one that would be required for the representations of abstract variables^38^, which allow for better generalization. In both cases the encoded variables are linearly separable, i.e., they can be decoded with a linear classifier. Also, in both cases, we varied the number of conditions corresponding to different combinations of values for the represented variables.

In most of the scenarios that we simulated the decoder performance is either disrupted or it remains the same when the correlations are destroyed. The beneficial effect of the correlations is maximal when the representations are fairly unstructured. In the case of random representations, the effect is maximal for a certain number of conditions. This number depends on the number of encoded variables and on the number of values that each variable can hold (i.e. the total number of conditions).

Our experimental observations that show that the performance of decoder is disrupted more in CA1 than in DG would be compatible with a scenario in which the representations in CA1 are unstructured, similarly to the simulated representations obtained with the random model. Our results also show that the representations in DG are compatible with at least two scenarios: 1) they could also be unstructured, but with a different number of encoded variables, either very small or very large, or, 2) the DG representations could be more similar to the more structured representations that we considered in Figure S15d-f. It is important to stress that the scenarios studied in Fig. S15 are all reasonable in the sense that they are based on representations that have already been observed in other studies^1, 38^. However, they are certainly not exhaustive and there could be other reasonable codes that we did not consider.

Our simulations in which multiple variables are encoded are compatible with recent models of the hippocampus that emphasize its role in memory compression^39, 40^, and memory prediction^41–45^: for all these models the neural representations in the hippocampus are constructed by learning the statistics of the sensory experiences in order to generate a compressed representations of the memories to be stored, or, when focused on temporal sequences, to generate a prediction of the next memory (successor representation). Future theoretical work will establish more quantitatively whether this scenario is fully compatible with our observations and what the different roles of CA1 and DG could be in this compression process.

All these results strengthen the hypothesis that the neural code in the DG and in the CA1 area of the hippocampus is highly distributed and that it is important to analyse it using a population approach^7, 18, 46^.

The analysis of the averaged response properties of individual neurons is certainly informative but it is not sufficient to characterize the neural code of a brain area. Critically, the role of the DG and the CA1 area of the hippocampus should be revisited in light of our observations. The methods that we propose will shed new light on the general role of other brain areas, in which place cells are not observed, in high level cognitive functions such as spatial navigation.

## Methods

### Calcium imaging

Mice were prepared for in vivo calcium imaging as previously described^15^. For dorsal DG imaging, mice were injected with a virus encoding GCaMP6m (AAVdj-CaMKII-GCaMP6m, (AAVdj-CaMKII-GCaMP6m, ∼4X1012vg/ml, Stanford Vector Core) at the following coordinates: − 1.95AP, 1.4ML, 2.2, 2.1, 2.0, 1.9 DV, ∼ 90nl per site) and a ∼ 1.0mm diameter, ∼ 4mm long GRIN lens (Inscopix, Palo Alto, CA) was implanted at (−2.0AP, −1.4ML, −1.95 DV). For dorsal CA1 imaging, mice were injected with a virus encoding GCaMP6f (AAV1-Syn-GCaMP6f.WPRE.SV40, U Penn Vector Core) at the following coordinates: (−2.15AP, 1.85ML, −1.55, −1.65DV, 256nl per site) and a GRIN lens was implanted at (−2.15AP, 1.30ML, −1.30DV). Three weeks after surgery, mice were checked for GCaMP expression with a miniaturized microscope (Inscopix, Palo Alto, CA) with procedures previously described^15^. Anesthetized mice were checked for GCaMP+ neurons and a baseplate was attached to the skull at the optimal imaging plane. For dorsal DG imaging, one week later, mice were imaged during foraging in an open field task and were habituated to the room and enclosure (30min), then 24 hours later they were imaged as they foraged for sucrose pellets in an open field enclosure (50cm^2^). For dorsal C1 imaging, mice were imaged during exploration of an open field enclosure. Mice were habituaded to the room and enclosure (10 minutes) and then imaged 30 minutes later. Imaging frames were recorded with nVista acquisition software (Inscopix, Palo Alto, CA), and time-synced behaviour was acquired using EthoVision XT 10. Calcium imaging videos were acquired at 15 frames per second with 66.56 ms exposure.

### Behaviour data pre-processing

The behaviour was recorded using a webcam (Logitech) mounted on the ceiling about 3 feet above the arena. The instantaneous position of the animal was then extrapolated from the video using custom code written in Python using the Scikit-image library (version 0.13.0). We first applied a 9 points piecewise affine transformation to correct for barrel camera distortions. We then applied a smoothing filter with a Gaussian profile to reduce the effect of pixel intensity noise due to low lighting and low image resolution and applied a threshold to the gray-scale converted image to get a few contiguous regions of pixels as candidate animal tracking. We then used a method based on the determinant of the Hessian to identify blobs in the pre-processed images and verified that the largest blob was consistently found to be corresponding to the animal silhouette. Hence, we used the centre of the largest blob as the tracked position of the mouse. We further temporally aligned the position data to the imaging data using linear interpolation and smoothed them with a 7 frames time window. Lastly, we identified the time bins in which the speed of the animal was lower than 2 cm/s for more than 1 s and discarded them from the analysis, unless specified.

### Signal extraction and spike deconvolution

All calcium movies were initially processed in Mosaic (Inscopix, Palo Alto, CA) for spatial binning and motion correction and subsequently analysed using a recently developed software algorithm written in Matlab (Mathworks) called CNMF-e^16^. Briefly, the algorithm separates the large, low-frequency fluctuating background components from the signal produced by of multiple sources in the data, allowing the accurate source extraction of cellular signals. It involves a constrained non-negative matrix factorization problem optimized for endoscopic data whereby calcium temporal dynamics and the shape of spatial footprints are used as constraints. It includes 3 main steps which are iterated: obtain a first estimate of spatial and temporal components of single neurons without direct estimation of the background; estimate the background given the estimated neurons’ spatiotemporal activity; update the spatial and temporal components of all neurons while fixing the estimated background fluctuations. In each of these steps, manual intervention guided by visual inspection based on temporal profile and spatial footprint shape allowed to further improve the quality of the signal extraction. The result of this process consists of a list of deconvolved calcium events for each cell with associated timestamp and magnitude and the convolved trace obtained with the estimated calcium decay profile. For our analysis, we used an exponential profile to integrate the calcium events in time with a time scale that optimized the position decoding performance, although we empirically observed that results did not qualitatively change across a wide range of reasonable integration time values (see Fig. S1).

### Place fields and heading direction tuning

Place fields for each extracted source were constructed in a manner similar to established method applied to electrophysiology data^10^. We used the calcium events of each cell as its putative spiking activity. We then summed the total number of events that occurred in a given location, divided by the amount of time the animal spent in the location and smoothed using a Gaussian kernel centered on each bin. The rate in each location x was estimated as

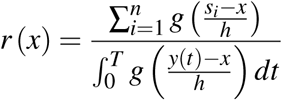

where *g* is a gaussian smoothing kernel, *h* = 5 sets the spatial scale for smoothing, *n* is the number of events, *s_i_* is the location of the *i*-th event, *y* (*t*) the location of the animal at time *t* and [0, *T*) the period of the recording. In this and all subsequent analysis we removed the time bins in which the animal had a speed of less than 2 cm/s for more than 1 s, unless specified otherwise. Similarly, for heading direction tuning, we first discretized the directions of motion into 8 angular bins of 45 degrees each and then computed the mean event rate for each cell in each of the 8 bins.

### Spatial information statistics

To quantify the statistical significance of the rate maps we measured their specificity in terms of the information content of cell activity^27^. We used a 16×16 square grid and computed the amount of Shannon information that a single event conveys about the animal’s location. The spatial information content of cell discharge was calculated as a mutual information score between event occurrence per cell and animal position or equivalently using the formula:

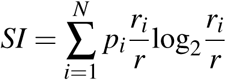

where *i* is the spatial bin number, *p* is the probability for occupancy of bin *i*, *r_i_* is the mean event rate at bin *i* and *r* is the overall mean event rate. We applied the same formula to the direction of motion after discretizing the full angle to 8 bins of 45 degrees. For both measures, we corrected for the sampling bias problem in information measures^28^ using shuffled distributions of event occurrences. We assigned randomly chosen time bins to each calcium event of a cell and computed the resulting information content. We repeated this operation for 1000 similar shuffled data for each cell independently. We then compared the original information content to the resulting distribution and considered a cell as a place cell or a head direction cell if the original value exceeded 3 sigmas from the shuffled distribution.

### Decoding position

For all the datasets, unless otherwise specified, we used 10-fold cross validation to validate the performance of the decoders. We divided the trial in 10 temporally contiguous periods of equal size in terms of number of datapoints. We then trained the decoders using the data from 9 of them and tested on the remaining data. To decode the position of the animal, we first divided the arena into 8×8 equally sized, squared locations. We then assigned at each time bin the label of the discrete location in which the animal was found. For each pair of locations, we trained a Support Vector Machine (SVM) classifier^47^ with a linear kernel to classify the cell activities into either of the two assigned locations using all the identified cells unless specified otherwise. We used only the data corresponding to the two assigned locations and to correct for unbalanced data due to inhomogeneous exploration of the arena we balanced the classes with weights inversely proportional to the class frequencies^48^. The output of the classifiers was then combined to identify the location with the largest number of votes as the most likely location^22^. The decoding error reported corresponds to the median physical distance between the centre of this location and the actual position of the mouse in each time bin of the test set, unless otherwise specified. To assess the statistical significance of the decoder, we computed chance distributions of decoding error using shuffled distributions of calcium event occurrences. Briefly, for each shuffling, we assigned a random time bin to each calcium event for each cell independently while maintaining the overall density of calcium events across all cells, i.e., by choosing only time bins in which there where calcium events in the original data and keeping the same number and magnitude of the events in each time bin. This method destroys spatial information as well as temporal correlations but I keeps the overall activity across cells. We trained one decoder on each shuffled distribution and pooled all the errors obtained. We finally assessed the statistical significance of the decoding error for the 10-fold cross-validation of the original data by comparing them to the distribution of errors obtained from the shuffled data using the non-parametric Mann-Whitney U test, from which we obtained a p-value of significance. We applied other strategies for assessing the significance of the decoder to verify that the results did not depend on the particular strategy adopted (see Fig. S3). We also report a comparison between different decoding strategies, including probabilistic approaches such as Näive Bayes, in the Supplemental Material (see Fig. S4).

### Decoding of the direction of motion

One behaviourally relevant quantity that was available to us was the direction of motion of the animal. Unfortunately, the visual tracking didn’t allow for a direct estimate of the direction of motion. The head direction was also not easily measurerable available so we resorted to using the positional information to extract the direction of motion. We computed it by using two subsequent datapoints in the animal x-y trajectory. We discretized the values into 8 angles and then applied similar decoding strategies as for position decoding, i.e., we used a battery of linear-kernel SVM decoders to distinguish between pairs of angles after balancing the dataset through class weighting. We report the median error in radiant on the left-out data of the 10-fold cross validation. We applied the methods described above for position decoding for assessing the statistical significance of the results.

### Decoding of speed

To decode the speed of movement of the animal we first computed the speed of motion using two consecutive positions and assigned the computed speed to the later time bin among the two. To decode the instantaneous speed of motion we used Lasso^49^, a linear regression analysis method that minimizes the sum of squared errors while selecting a subset of the input cells to improve decoding accuracy and interpretability of the results. For assessing speed decoding chance level, we applied the same strategy as for the other variables except we shuffled all the events of each cells in time independently and convolved them with the same temporal profile we used for the original data.

### Importance index

The importance index was introduced to quantify the contribution of each cell in a population to the decoding of a given quantity. We applied a modified version of a traditional method for feature selection in machine learning. In our analysis, a feature of the input space consists of one DG cell. Feature selection is performed using the weights of the decoder after fitting model to the data. In our case, since we employed multiple decoders, one for each pair of physical location in the arena, we introduced a method to combine the weights assigned to the cells by each decoder. We defined the importance index of cell *i* as:

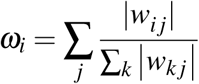

where *w_i j_* is the weight of the *j*-th decoder assigned to the *i*-th cell.

### Procedure to destroy correlations

To destroy correlations without impacting the spatial information of single neurons, we considered multiple passes through single discrete locations in the arena. We then shuffled the calcium event occurrences between different passes in the same location. Importantly, we corrected the activity of each pass for the different amount of time spent in each pass by radomly sampling events instead of replacing them in order to reduce artifacts. We verified that the correction does not impact decoding by sampling from the same pass (see Fig. S14).

### Code and data availability

The data analysis has been performed using custom code written in Python (version 2.7.12) and routines from the Scipy (ver. 0.19.0), Numpy (ver. 1.11.3) and the Scikit-learn (0.19.1)^48^ packages. The source extraction has been performed using Matlab (Mathworks, R2016a) and an early version of the CNMF-e^16^ package. The source code and the data are made available upon request.

## Acknowledgments

This article is dedicated to Howard Eichenbaum. We thank Loren Frank, Surya Ganguli, Liam Paninski, Daniel Salzman, Mattia Rigotti and Marcus Benna for insightful discussions and comments on an earlier version of this work and Pengcheng Zhou for his help with the CNMF-e software. FS was supported by the Swiss National Science Foundation’s Early Postdoc. Mobility grant P2EZP2 155561 and by the Kavli Foundation. SF was supported by NSF’s NeuroNex program award DBI-1707398, the Gatsby Charitable Foundation, the Simons Foundation, the Schwartz foundation, the Kavli foundation. MK was supported by NIMH R01 MH108623, R01 MH111754, and IMHRO/One Mind Rising Star Award. JHJ was supported by the Helen Hay Whitney Foundation.

