## Supplemental Material for "A distributed neural code in the dentate gyrus and in CA1"

### Fig. 1: Preprocessing; Related to Fig. 1

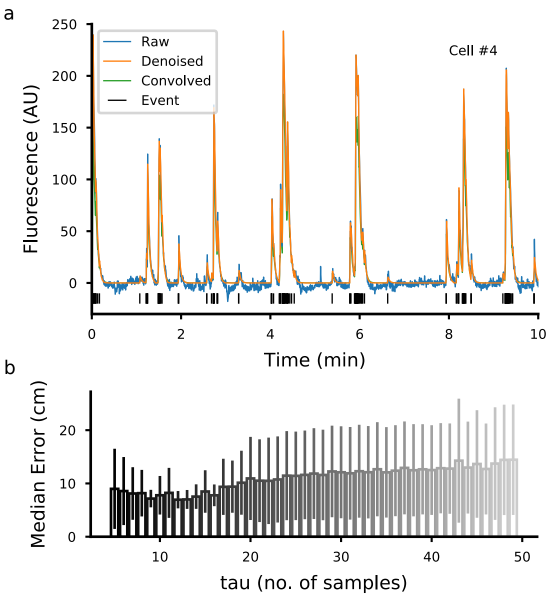


Here we illustrate the type of signals extracted using CNMF-e that we used for our analysis. The algorithm's output is composed of a spatial footprint, the raw calcium signal, the denoised signal and the calcium events timings and magnitude for each cell. The raw signal corresponds to the fluorescence levels attributed to the single sources when the contaminating signal due to neuropil in the background and to neighbouring sources have been eliminated. This signal includes noise due to the recording apparatus. The denoised signal is obtained using an auto-regressive model that uses the estimated calcium event timings. The parameter of the model as well as the calcium event times, the spatial footprints and the background estimation are all part of the minimization process in CNMF-e [16].

In our analysis we focused on the calcium events times because these gave us the closest estimates of each cell's activity. We then convolved the calcium events in time with a decaying temporal profile to cumulate information in time. Similar procedures have been used in the past to amplify the signal with temporally sparse data [17].

**a**) Example of fluorescence signal extracted using CNMF-e [16]. The raw, denoised and event signals correspond to the output of the algorithm. The convolved signal is the one used in this work and is obtained by convolving the events with an exponential kernel of a duration that optimizes the position decoding performance (see **b**).

**b**) Median Error (mean ± s.d.) of the position decoding for different time scales of the exponential kernel used to convolve the calcium events. The chosen time scale corresponds to the one that minimizes the position decoding error (2.2 s in this example, 12 samples).

### Fig. 2: Place-cells; Related to Fig. 4

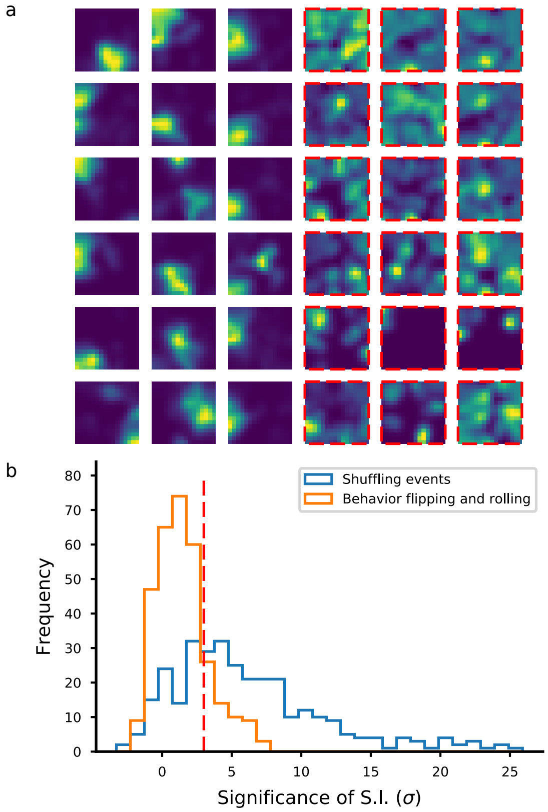


In order to quantify the spatial tuning of the cells, we applied standard methods used in electrophysiology to measure the spatial information contained in the activity of each cell using the calcium events [10]. We corrected for the sampling bias problem [7] by using a shuffling approach. We quantified the tuning of the cells by the statistical significance of the spatial information content (SSI). Similar to what we did for assessing the chance level decoding (see Fig. S), we used two shuffling methods for assessing the SSI of each cell. The first method (blue histogram) consisted in shuffling the event timings, as it is commonly used in the definition of a place cell. We identified place cells as those with an SSI>3. For comparison, here we report a second method which consisted in time reversing the animal's trajectory and then shifting it in time by a random amount. The second method preserves the activity of all cells but and it only destroy its association to the position of the animal and therefore it's more conservative.

**a**) Gaussian-smoothed event density maps for a selection of cells, normalized by the mouse's occupancy time per 16x16 unit area and the cell's maximum response in the 50 cm x 50 cm arena [17]. The cells have been ordered by the statistical significance of their spatial information, from the most to the least significant. Here we show the group of 18 cells with the most significant SSI (place cells) and the 18 with the least significant SSI (non place-cells, red-borders) from the dentate gyrus of a representative mouse (DG3).

**b**) we show the histogram of SSIs for all the DG cells recorded for mouse DG3 computed with the event shuffling method (blue) and the trajectory shuffling method (orange). The latter method highlights the importance of adopting the correct null hypothesis to assess the spatial information of the cells. In our recordings, many of the cells that are classically classified as place cells have a spatial information content that is not statistically different from the one obtained by a conservative shuffling of the trajectory which still disrupts the association between a cell's activity and the mouse position at each time point. In the text, we used the former definition for consistency with the literature.

### Fig. 3: Fraction of place cells in DG and CA1; Related to Fig. 4

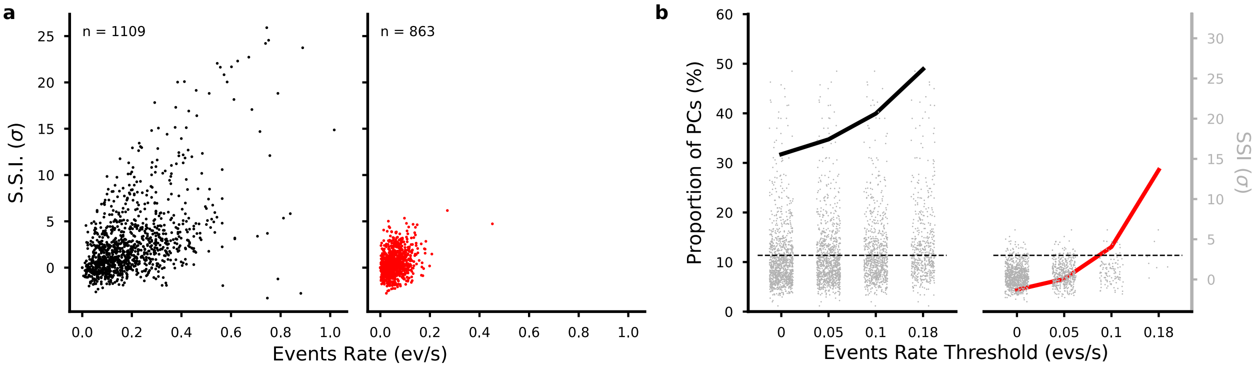


Reported values of place cells ratios in CA1 and DG vary considerably across labs. This is due to a combination of factors, including different recording techniques, different statistical methods used to assess spatial tuning and differences in experimental protocol. Our imaging data is unique in that a systematic comparison of spatial tuning of pyramidal cells CA1 and granule cells DG in a freely moving animal has never been reported so far. Here we compare the significance of spatial information in the populations of CA1 and DG cells we identified. In both areas, place cells were the minority of cells, however place cell ratios where very different (36% in DG, 4.2% in CA1). One reason for this difference could be in the lower activity levels we found in CA1 cells compared to DG. Place cell studies in rodents hippocampus have found higher place cell ratios when animals run through a linear track [17]. In [11] the authors found that 47.8% of the recorded CA1 cells were place cells but they used spatial information measure alone [10]. The lower ratio of place cells in our report could be due to a higher sensitivity for sparsely active cells of our method with respect to standard extracellular recording approaches. Despite the likely presence of some false negatives in the single calcium event detection, even cells with a very small number of sufficiently strong calcium events are detected by the source extraction algorithm we used. Also in other recent studies, Calcium imaging has revealed larger proportions of sparsely active cells suggesting these populations have been largely underestimated in electrphysiology studies [4, 12]. In particular, the CNMF-e algorithm uses both spatial footprints and temporal components to separate cells from background contamination and from other sources [16].

Instead, typical clustering methods for extracellular recordings may bias identification towards more densely active cells because these are more likely to be pulled out from the noise through clustering. This bias ultimately results in an overabundant representation of place cell ratios in the population.

To validate our hypothesis, we looked at the relation between activity levels in terms of event rates and spatial information (**a**). We removed weakly active cells in both CA1 and DG populations with a varying threshold to see if we could artificially bias either population to artificially high place cell ratios. This was indeed the case as we could obtain higher place-cells ratios, compatible with the literature on electrophisiology studies, by choosing an appropriate threshold on activity (**b**). This could mean that some CA1 cells are more sparse than previously thought and CA1 place cells may have been over represented in place cells studies.

**a**) Significance of spatial information (SSI) as a function of event rates across a session for all cell populations in DG (left) and CA1 mice (right).

**b**) Place cells ratios after removing cells based on their mean activity. For each threshold on activity, we report the SSI for the selected cells (gray dots) and the corresponding proportion of place cells in the remaining population. For example, if we didn't record any signal from cells with less than 0.18 events per second, we would have observed about 30% of place cells in CA1 and 50% of place cells in DG across all mice. The dashed line corresponds to the threshold on SSI used to classify cells as place cells.

### Fig. 4: Dependance of decoding performance from the number of cells in CA1 and DG; Related to Fig. 2

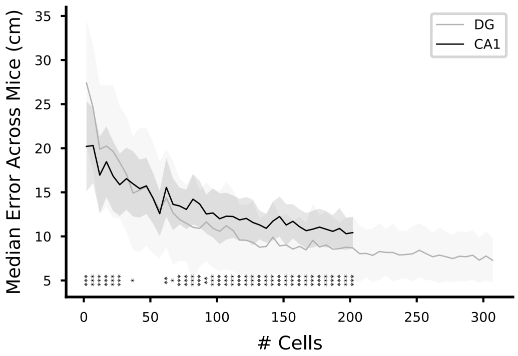


Decoding position performance using different number of cells. For each given number of cells, a random selection of cells from the pool of identified cells is used to decode the animal's position using 10-fold cross-validation. The selection is repeated 10 times for each animal and for each number of cells. We then pooled all the results from all the animals to report the mean (solid line) and standard deviations (shaded area) of the median errors for position decoding for each given number of cells (indipendent samples t-test for significance, $*p<0.05$, $**p<0.01$, $***p<0.001$). The maximum number of cells we used corresponds to the minimum number of cells we could record from any animal in DG or CA1.

### Fig. 5: Chance level decoding performance; Related to Fig. 2

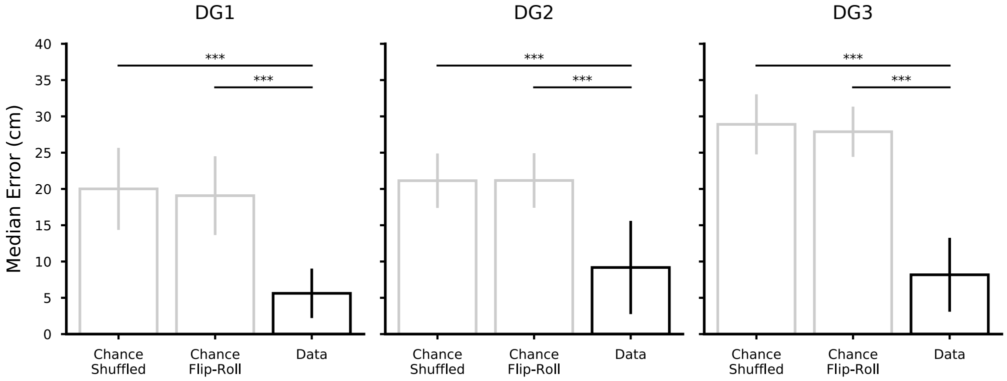


It is important to assess the performance of the decoder to a chance performance, i.e., the performance of a prediction that is not based on the recorded activities but solely on the behaviour of the mouse. The important factors that must be considered in the case of decoding continuous behaviour, are represented by the auto-correlations of the neural activities and of the behaviour. Therefore, for evaluating the chance performance one must take into account these factors in order to avoid underestimating the performance obtained by chance and therefore overestimate the ability of the decoder to extract the information under consideration.

We computed the chance performance by training a position decoder on artificially manipulated data. We used two methods: the first method consisted in shuffling the calcium events in time by keeping the overall instantaneous event rate across the population constant ('chance shuffled'). The second method consisted in applying a time reversal operation to the animal x-y trajectory and then shifting it in time by a random amount. Data points falling outside the temporal window due to the shifting where reinserted from the other end of the period as in a torus.

We then trained the decoder on a cross-validated manner in which the data was split into 10 chunks of contiguous data, 9 of which were used for training and the remaining one for testing the decoder. For both methods of shuffling, therefore, we obtained the 10-fold cross validated performance which we aggregated all together in each animal and applied a Mann-Whitney U non-parametric test to compare the resulting distribution to the original data (***$p<0.001$). In this figure we report the performance (mean $\pm$ s.d.) for the position decoders for all the animals for the two chance levels in grey and for the original data in black. The decoding performance is significantly above chance for both choices of chance level and the difference between the two methods is small.

### Fig. 6: Comparison of different decoding strategies; Related to Fig. 2

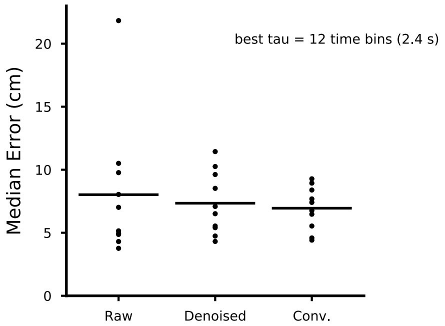


To decode the animal's position and direction of motion, we used a battery of linear decoders trained on pairs of locations (see Methods). In this way we could use the weights of the single decoders, properly combined, to obtain an overall importance index for the cells in the population. However, the animal's position could be decoded also by means of a more traditional approach using 'optimal decoding' [15]. In **a** we report the position decoding performance (mean $\pm$ s.d.) from the dentate gyrus of a representative animal (DG3) using both a battery of linear decoders (support-vector machines with a linear kernel, SVM) [3] and a näive Bayesian approach [14, 15]. The performance of the two types of decoders is not significantly different (Mann-Whitney U test, $p > 0.05$). The routines used come from the Python scientific package Scikit-learn [8].

Another important choice is related to the type of signals used for decoding. Using the convolved calcium events instead of the raw traces as described in Fig. S gives a minor (not significant) improvement on the decoding position performance (data for a representative mouse DG3 in **a**).

### Fig. 7: Position decoding error and speed are negatively correlated; Related to Fig. 2

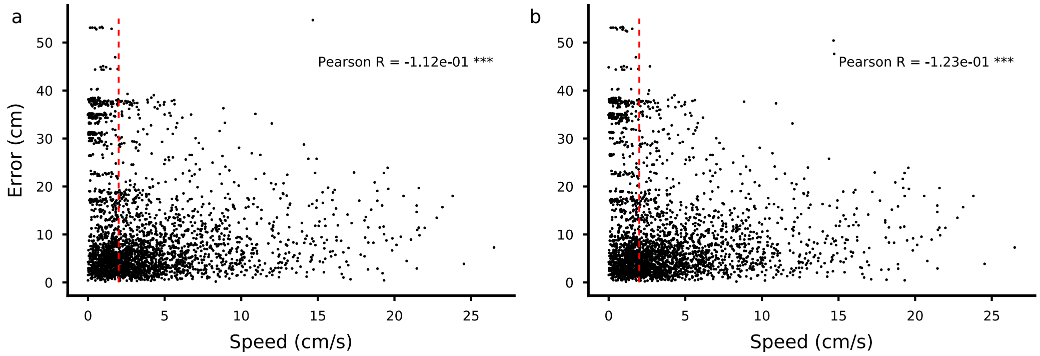


A mouse's movement and immobility states are often considered two distinct behavioral conditions, characterized by distinct neural activities, and have been therefore considered separately for analysis in the literature. In our work, we focused our analysis on the data collected during movement following a typical procedure in the hippocampal research, see for example [6]. We then asked whether the accuracy the decoded position was related to the speed of movement. First, we decoded the position of the animal using only the datapoints during movement for training and testing on all datapoints. In this figure we plot the instantaneous error on the test datapoints of the 10-fold cross-validation (see Methods). On the x-axis, we plot the instantaneous speed of movement in one animal (data from the dentate gyrus of a representative animal DG3). The decoded position is taken as the centre of the discrete location that is selected by the decoder in each time bin. The red vertical line corresponds to the value we used as a threshold to distinguish between movement and immobility on the training data. To verify that the negative correlation was not due to the fact that the datapoints corresponding to immobility were not used in the training set, we repeated the procedure using all the datapoints during training. The negative correlation was still present (Pearson $R = -0.123$, $^{***}p < 0.001$). This analysis confirmed that the position is encoded more accurately when the animal moves.

### Fig. 8: Temporal stability of the importance index; Related to Fig. 4

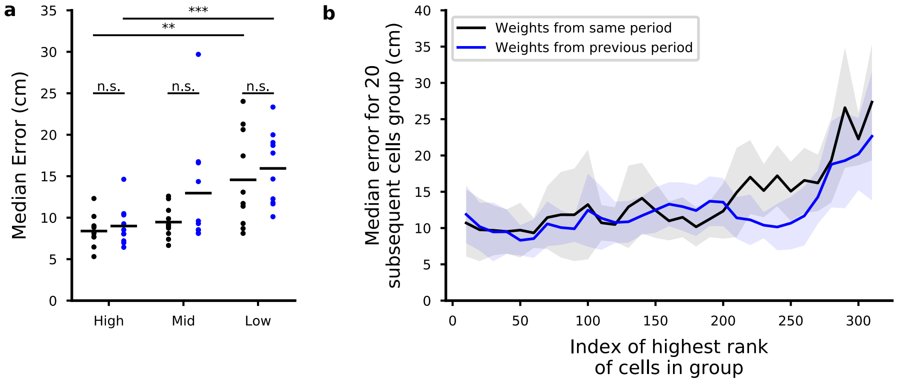


We introduced an importance index that quantifies the contribution of each cell to the population code. We know from the analysis in the main text that the ranking of the cells based on the importance index depends on the cells that are considered for decoding. So we know it is not an intrinsic property of the cell. However, given a population of cells that is used to estimate the importance index, it is interesting to ask whether the rank of a cell changes over time.

**a**) We ranked the cells from the dentate gyrus using the first 10 minutes of data from the 30 minutes trials. We computed the decoding performance using the 20 best cells (ranked high), 20 cells that ranked in the middle and the 20 cells in the lowest part of the ranking. We then decoded position using each group of cells. In particular, we decoded the position of the animal in the next 10 minutes using the same groups of cells. If these cells' ranking changed, we'd expect a different performance for each group of cells since they were defined using the ranking in the first 10 minutes. Instead, we see no significant changes in decoding performance for the hight, middle and low-ranked cells. This indicates that the importance of each cell is relatively stable over time, at least when the decoding performance is considered.

**b**) We then applied a procedure like that of Fig. 4 of the main text to better assess whether the ranking in one time-period could be used to rank cells in another time period. We first computed the decoding performance in the first 10 minutes of the trial using subsequent groups of cells ranked by their importance index computed in the same time window. We then compared these results with the ones obtained by decoding position during the next 10 minutes of the trial. The two plots largely overlaps, therefore the importance indices obtained in one temporal window can be used to select the important cells in another temporal window.

### Fig. 9: Importance index and spatial information; Related to Fig. 5

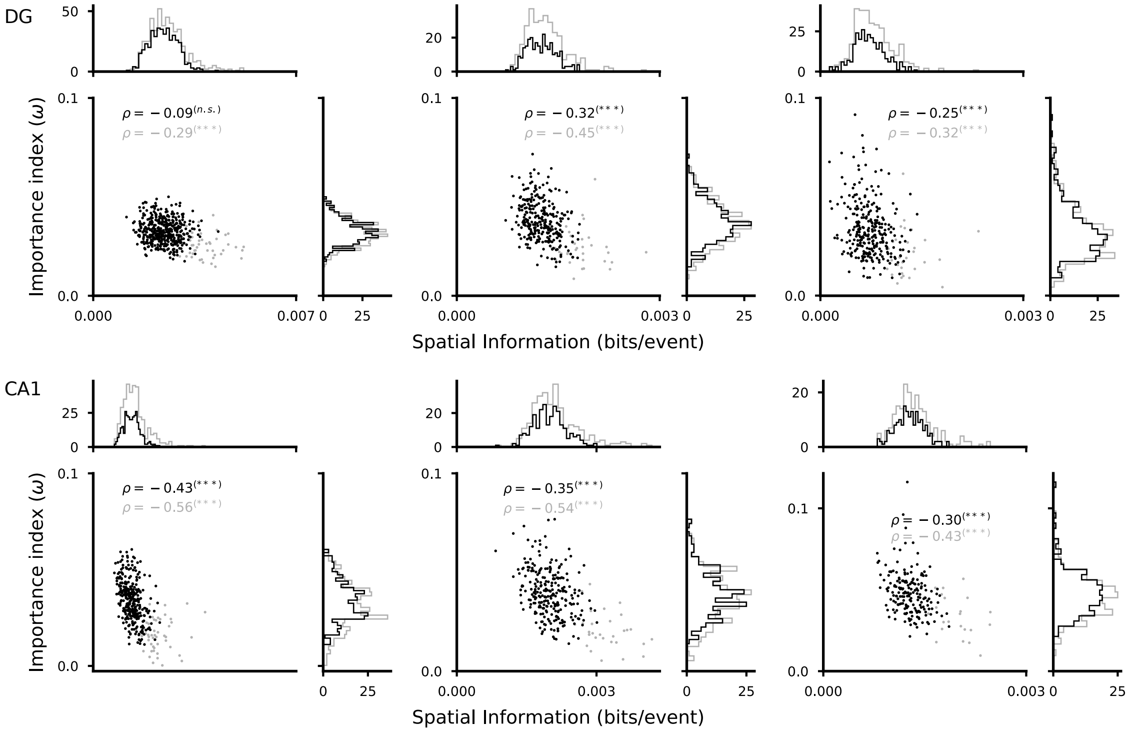


Limited data, such as low firing rate, introduce a systematic bias in the estimation of the information content in neural activities [7]. Our results show that a more sensible choice is the one of the SSI, a measure of the strength of the spatial information after compensating for this bias. Our results show that the importance index was weakly correlated to a measure of the strength of the spatial information contained in the cell activities, which we quantified using the SSI. The SSI is a sensible choice because of the systematic error in the estimation of the information content with limited data, as it is often the case with neural data [7]. We also showed that importance index and SSI are correlated. Here instead we further show that importance index and spatial information have a low and even negative correlation factor (Pearson-R correlation, $***p < 0.001$). In this figure we report the scatter plots of the importance index for position and spatial information for each DG and CA1 animal. Each dot in the plots represents one cell (grey: all cells, black: cells with more than 10 calcium events identified). The histograms for each of the quantities are shown on the sides of each plot. This results further validates the choice of the SSI over the one of raw spatial information as a measure of how much a cell encodes position in a given brain area.

### Fig. 11: Control for correlation between movement direction and position; Related to Fig. 7

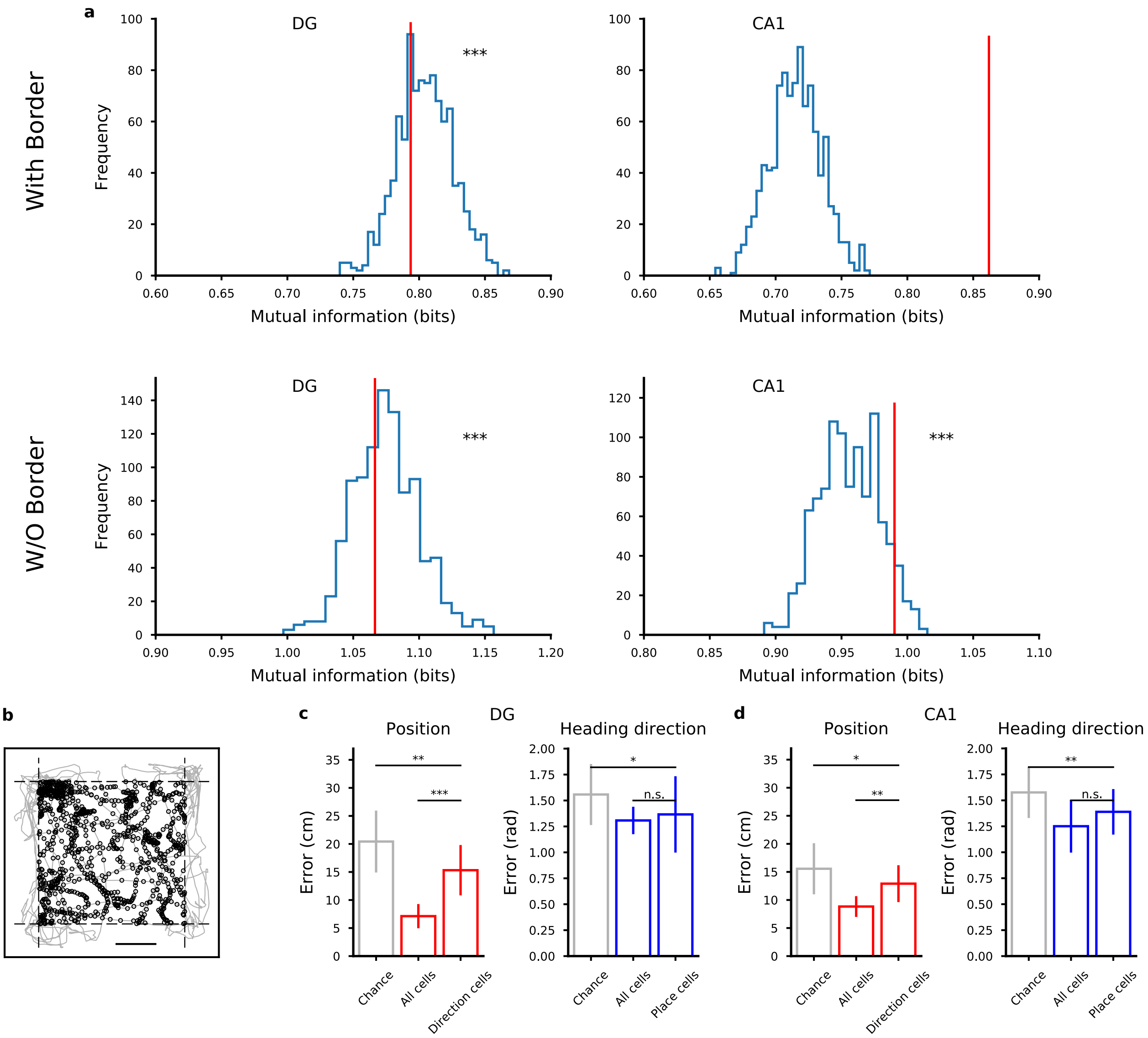


We showed in the main text that we can separately decode position and direction of motion and that the neural code for these two variables appears distributed across the neural population (Fig. 7). Our strategy was to show that the important cells for decoding position could be used to decode the direction of motion. However, it could be that the reason why direction could be decoded using the important cells for position was that direction and position were correlated. Indeed, when the animal is very close to one the border of the arena, the direction of movement can only assume values that point towards the inside of it. The decoder could in principle use this information to decode direction of motion from the important cells for position, therefore confounding our results.

To exclude this possibility, we computed the mutual information between direction of movement and position and compared it to the distribution obtained with 500 shufflings of data from the animal with the most homogeneous coverage of the environment (**a**, DG in top left panel, CA1 in top right panel, same animals as in Fig. 7). In the shuffled data, position was time reversed and shifted similarly to what was done for evaluating the chance performance (see also Methods in the main text). The mutual information in the real data (red line) lies within the distribution of mutual information for shuffled data (blue histogram, one sample t-test, $^{***}p < 0.001$) for DG, hence the knowledge of one variable is not enough to predict the value of the other variable, but not for CA1. We attribute this difference to the effect of the borders in the CA1 experiments, in which a rectangular arena of half the size than the one for DG mice was used. Hence, to further verify that the correlation between direction of motion and position did not explain the results of Fig. 7b, we repeated the decoding analysis after removing from the data all the positions that were recorded in the spatial bins closest to the walls of the arena (**b**, example DG mice). After this manipulation, the mutual information between position and head direction becomes indistinguishible from the chance distribution in both DG and CA1 (bottom panels in **a**) and we were still able to decode position from the most important cells for direction and vice versa, as we did for the original data in the text, for both DG (**c**) and CA1 mice (**d**).

**a**) Mutual information between position and direction of motion for shuffled data (blue histogram) and real data (red vertical line) for original data with positions close to the arena walls and for filtered data as in **b**.

**b**) Positions close to the walls of the arena, i.e., beyond the dashed lines, were eliminated from the data. Grey: mouse trajectory. Black dots: positions included in the analysis. Same DG mouse as in Fig. 7.

**c**) Decoding error for important cells for position (left) and direction (right) from a representative DG mouse, same procedure as in Fig. 7b in the main text but after removing border data as in b.

**d**) Same as in **c** but for the CA1 mouse of Fig. 7b.

### Fig. 12: Temporal stability of the decoder; Related to Fig. 2

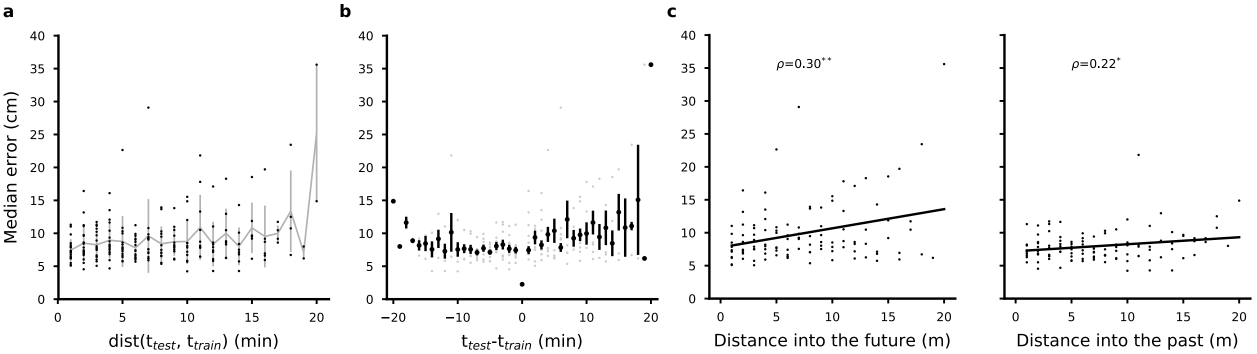


We asked whether the representations for position changed within a recording session. To do this, we used the 30 minutes trial available in our data (animal DG3, data from the dentate gyrus). We trained a position decoder on a shifting window of 10 minutes of data and tested in on a shifting window of 1 minute of data. The performance of the decoder is remarkably strong and stable for a period of time extending to the available 20 minutes in the trial, although with some slow degradation towards the more distant time windows (**a-c**). This result shows that position is encoded with strong accuracy on the populations of cells whose coding properties extend long in the trial and it seems compatible with results in literature showing that the representation of position is stable across session despite the large degree of variability on the tuning properties of single cells [17].

We further looked for evidence of a difference between testing the decoder in past time periods and future time periods with respect to the training period. To determine whether decoding data temporally preceding the training data (past) was different from decoding following data (future), we linearly interpolated the decoding performance and obtained two weakly significant Pearson correlation factors of 0.3 for the future ($^{**}p < 0.01$) and 0.22 for the past ($^{*}p < 0.05$). Given the weakness of our statistical test we couldn't detect a difference between these two trends. Therefore, the performance on the test data does not depend on whether the decoder was trained on a past or on a future interval.

**a**) Position decoding performance as a function of the distance in time between training set and test set. Each dot corresponds to one choice of training and test set, the grey line joins the mean values on pooled data for each bin of distance in minutes, bars are standard deviations within the binned distances.

**b**) Data as in (**a**) but datapoints for which the test precedes the training set are on the negative side of the x-axis.

**c**) Data as in (**b**) where the points have been fitted with a linear regression for test data that is in the future with respect to training data (left) or in the past (right). Reported correlation values correspond to the Pearson's R correlation (*p<0.05, **p<0.01).

### Fig. 13: The information about position is distributed: distributions of importance indices in DG and in CA1; Related to Fig. 5

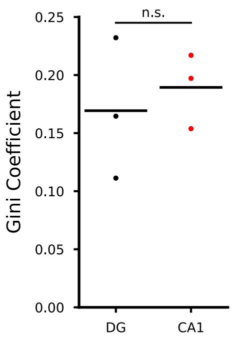


One way to compare how populations of cells represent information is to compare the distribution of their importance indices.

We considered the Gini coefficient [5], a measure of statistical dispersion among values of a certain frequency distribution. It is often used as a measure of inequality of income levels. Low values correspond to a large degree of equality, i.e., people share large proportions of the total wealth of the population. High values often identify situations in which wealth is concentrated on a small number of people. We used the Gini coefficient to assess whether the information about position is distributed across multiple cells, or it is carried by a small minority. We found very low Gini coefficients in all DG and CA1 mice, with a slightly higher mean value for CA1 mice. To put numbers into context, the values we found were lower than the one of the country with the lowest inequality in the world and less then half than the income inequality in the United States of America as of 2017 (0.41) [13].

### Fig. 14: Procedure to destroy correlations; Related to Fig. 8

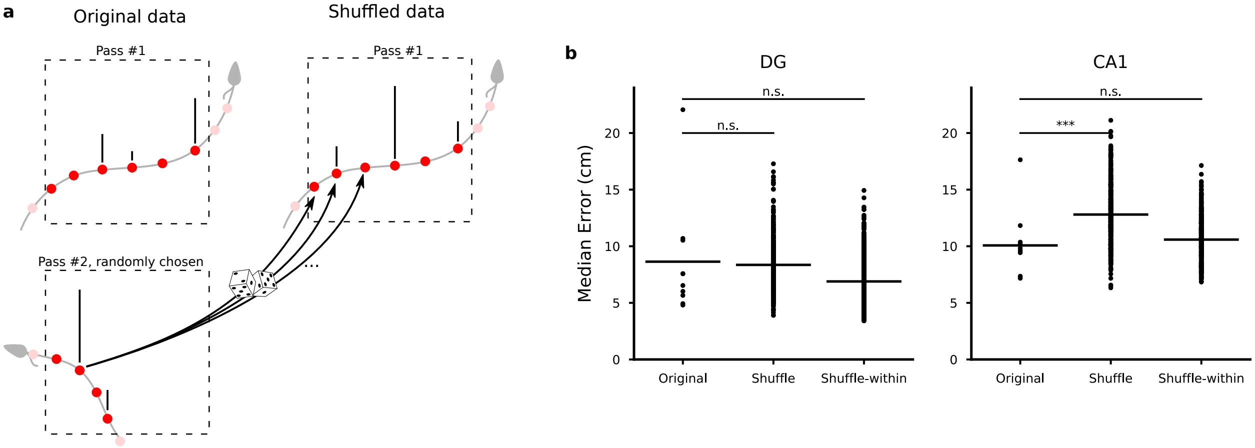


We devised a shuffling procedure to destroy correlations among neurons without affecting their spatial tuning profiles (**a**). For each location, we focused on all the passes of the animal through that location and identified the entrance and exit times of each one. Then for each pass, we substituted the recorded calcium events in that pass with the ones of an other randomly chosen pass. To account for the different number of time bins in the different passes, we randomly sampled from activity from the second pass. This procedure showed an effect in CA1 data but not in DG (**b**, one representative mouse for each area), suggesting different coding properties as discussed in the text. To verify that random sampling did not introduce artifacts that would affect decoding performance, we verified that when we sampled from data within the same pass the decoding performance was not significantly affected (**b**,'shuffle-within'). After this procedure, we were confident that any effect we would observe was only due to having destroyed correlations among neurons. When we shuffled data within the same pass, we saw no effects in either CA1 nor DG data.

**a**) Schematic description of the procedure for destroying correlations in one discrete location in the arena. The red dots correspond to the time bins of the recorded calcium events along the mouse trajectory (gray line) through the area of interest (dashed line). Time bins in which an event was recorded are identified with a tick mark on top of the red mark. Light red marks correspond to datapoints recorded before the animal enters and after it exits the area and are not considered. For each pass (pass 1 in the figure) we randomly select a second pass. We then generated shuffled data for each time bin of pass 1 by randomly sampling event data from pass 2.

**b**) Decoding error for original data, shuffled data as in a and shuffled data when the new data was generated from the same pass to control for artifacts due to sampling. Left, DG. Right, CA1 (Mann-Whitney U test, ***$p<0.001$).

**Fig. 15: The effects of dimensionality on neural correlations and decoding performance; Related to Fig. 8**

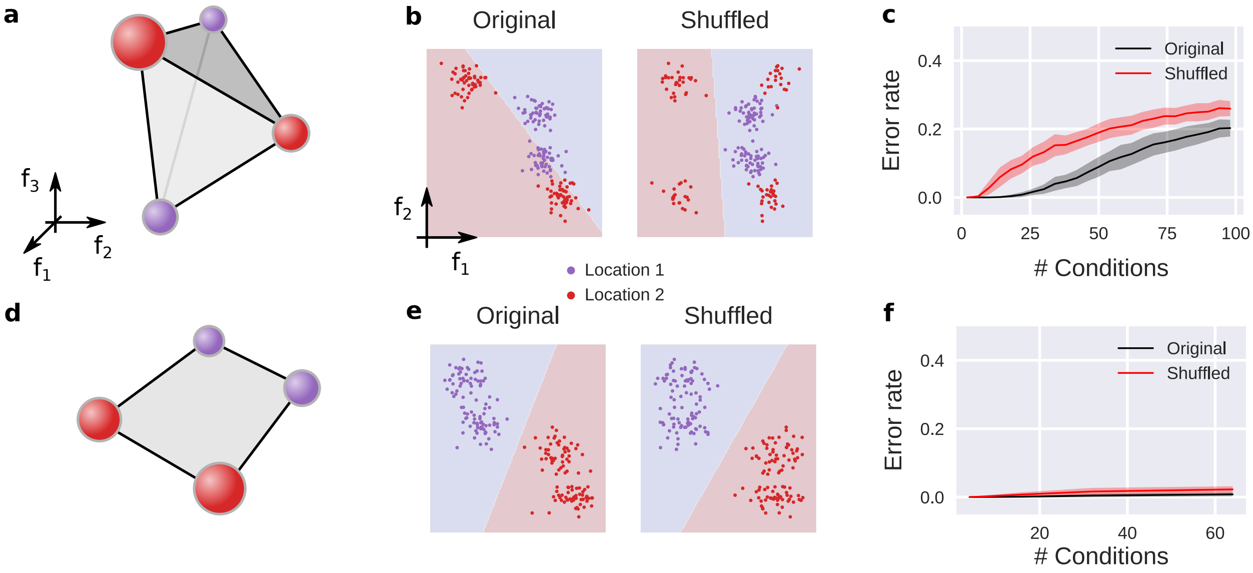


In our work we suggested that, at least in CA1, the neural correlations should not be considered as noise but rather as a reliable signal. We showed that destroying the neural correlations had an effect on the ability of our decoder to predict the animal position from CA1 activities. The effect was smaller in DG. In general, the effect of destroying correlations on decoding performance depends on the geometry of the neural representations [1], i.e., how the patterns corresponding to the different combinations of behavioral variables are distributed in the space of population activities. How do the effects of destroying correlation depend on the geometry of the neural representations? Here we studied in simulations a few cases to understand under what assumptions we should expect a disruption of the performance when the correlations are destroyed.

We started by considering neural representations in the space of population activities. These are determined by different combinations of discrete behavioral variables, each of which corresponds to an experimental condition. In the experiments, some of the behavioral variables would be under direct control, such as the discretized animal's position or movement direction, while others are not. For each condition we generated a prototype pattern of population activity and a cloud of points around it by adding isotropic Gaussian noise of zero mean and unit variance (see ). This cloud of points thus represents a set of recordings for that condition, similar to the ones generated by multiple passes of the animal through a given location in the environment as in Fig. 3 in the main text. Following our hypothesis, we generated many such clouds in the space of neural activities for the various combinations of hypothethical behavioral variables the neural activity is subject to and assigned half of them to one class, corresponding for example to one location, and half of them to another class, corresponding to a second location. The fact that the representations of each location consists of multiple conditions, naturally induces correlations among neurons in the population. This is because the activity of one neuron is tightly coupled to the activity of other neurons as prescribed by the patterns corresponding to that location.

To study the effects of destroying correlations between, we first considered the cross-validated ability of a linear decoder to correctly classify the two hypothetical locations. Then, we shuffled the data in a way similar to what we described in the Methods to destroy the correlations among neurons while keeping the information on location. Briefly, for each location, we chose one observed level of activation in that location for each neuron independently. This effectively violates the coupling imposed by the patterns of the conditions representing one location and therefore destroys correlations among neurons. Importantly, while this manipulation effectively destroys the correlations among neurons, it does not affect the statistics of activations for each neuron with respect to each location, i.e., the spatial tuning of the neurons is not affected. We repeated this procedure multiple times to generate a new dataset of data without neural correlations.

After this manipulation we trained again our decoder and compared the decoding performance with the one computed on original data, as we did for the neural recordings in the main text.

To being our study of how the geometry of the neural correlations affects decoding, we considered two common scenarios observed in neural data [2, 9], one in which the neural representations are 'unstructured' and one in which they are 'structured'. The first situation can be generated by randomly distributing the different conditions in the space of neural activities, as in **a**. For the second situation we considered abstract variables, i.e., the conditions were positioned at the corners of a low-dimensional hypercube randomly rotated in the space of neural activities as in **d** [2]. To visually describe the situation, in **b** and **e** we show these two scenarios in the simple case of two neurons. We further restrict our description to the case of two conditions per location, which could correspond for instance to two directions of motion in the data. The colored regions separated by a straight line represent the decision function of a linear classifier trained on cross-validated data, hence the red points lying on the red regions are correctly classified as are the purple points lying on the blue region. Vice versa, the points lying on a region of different color are incorrectly classified. In the case of unstructured data, the four clouds of points appear randomly distributed (**b**) and, in this example, they are linearly separable. The multiple conditions impose correlations between the two neurons, in a way similar to what we discussed in Fig. 3 in the main text. Indeed, after destroying the correlations, the two location are no longer linearly separable and therefore the performance of the decoder can only worsen. The situation in the high-dimensional space of the population activities doesn't have such a straight forward graphical interpretation and so it is not straight forward to predict what would happen in the high-dimensional case. Thus, we performed simulations in the high-dimensional case (50 neurons) and compared the decoder performance before and after destroying the correlations for increasing number of conditions (**c**). We found that the effect of the correlations increases with the number of conditions (**c**), it reaches a maximum and then it decreases again for higher numbers of conditions. Moreover, when the number of conditions becomes comparable to the maximum number of neurons (50 in our simulations), the two locations may be non-linearly separable also before destroying correlations, depending on the specific random arrangement, and so the error rate increases also for the original data.

In the case of structured neural representations, the situation is different (**d**, **e**, **f**). For this scenario, we generated the patterns in a way that follows abstraction as defined in [2] (**d**). As in the previous case, we first looked at a low-dimensional case of two neurons and two conditions per location (**e**). In this case the effect of the correlations is much weaker, as it is shown by the fact that also after destroying correlations the representations of the two locations are still linearly separable. To numerically explore the high-dimensional case, we performed similar simulations as before and varied the number of conditions while preserving the structure of abstraction (**e**). We observed that even for a large number of encoded variables the effect of destroying correlations was negligible in this case.

Taken together, these results show that the geometry of the neural representations imposes neural correlations can be beneficial for decoding performance. In some of the cases we examined we also found that the effect can be negligible. It is important to note that there can be other scenarios that we did not consider here but even with the few simple cases we described we found several conditions that seem compatible with our experimental data. Furthermore, it is interesting to note that the type of neural correlations that we considered in these models are not induced by a non-negligible covariance in the direction of the noise of neural activities. Rather they are induced by the presence of relatively small regions of the space of population activities that correspond to reliable representations to experimental conditions. It will the subject of further studies to investigate what classes of models can lead to such sparse representations.

[2] Silvia Bernardi, Marcus K Benna, Mattia Rigotti, Jerome Munuera, Stefano Fusi, and Daniel Salzman. The geometry of abstraction in hippocampus and prefrontal cortex. *bioRxiv*, page 408633, 2018.

[3] C Cortes and V Vapnik. Support-vector networks. *Machine learning*, 20(3):273–297, 1995.

[4] Mario Dipoppa, Adam Ranson, Michael Krumin, Marius Pachitariu, Matteo Carandini, and Kenneth D Harris. Vision and locomotion shape the interactions between neuron types in mouse visual cortex. *Neuron*, 98(3):602–615, 2018.

[5] Corrado Gini. Variabilità e mutabilità. *Reprinted in Memorie di metodologica statistica (Ed. Pizetti E, Salvemini, T). Rome: Libreria Eredi Virgilio Veschi*, 1912.
